# The homeodomain transcriptional regulator DVE-1 directs a program for synapse elimination during circuit remodeling

**DOI:** 10.1101/2022.10.21.512874

**Authors:** Kellianne D Alexander, Shankar Ramachandran, Kasturi Biswas, Christopher M Lambert, Julia Russell, Devyn B Oliver, William Armstrong, Monika Rettler, Maria Doitsidou, Claire Bénard, Michael M Francis

## Abstract

An important step in brain development is the remodeling of juvenile neural circuits to establish mature connectivity. The elimination of juvenile synapses is a critical step in this process; however, the molecular mechanisms directing synapse elimination activities and their timing are not fully understood. We identify here a conserved transcriptional regulator, DVE-1, that shares homology with mammalian special AT-rich sequence-binding (SATB) family members and directs the elimination of juvenile synaptic inputs onto remodeling *C. elegans* GABAergic neurons. Dorsally localized juvenile acetylcholine receptor clusters and apposing presynaptic sites are eliminated during maturation of wild type GABAergic neurons but persist into adulthood in *dve-1* mutants. The persistence of juvenile synapses in *dve-1* mutants does not impede synaptic growth during GABAergic remodeling and therefore produces heightened motor connectivity and a turning bias during movement. DVE-1 is localized to GABAergic nuclei prior to and during remodeling and DVE-1 nuclear localization is required for synapse elimination to proceed, consistent with DVE-1’s function as a transcriptional regulator. Pathway analysis of DVE-1 targets and proteasome inhibitor experiments implicate transcriptional control of the ubiquitin-proteasome system in synapse elimination. Together, our findings demonstrate a new role for a SATB family member in the control of synapse elimination during circuit remodeling through transcriptional regulation of ubiquitin-proteasome signaling.

**Contributions Summary:** KDA generated strains, transgenic lines, molecular constructs, confocal microscopy images and analysis, performed optogenetic behavioral experiments, photoconversion experiments, modencode ChIP-seq analysis and pathway analysis. SR performed all calcium imaging experiments/analysis and conducted single worm tracking. KB performed all Bortezomib inhibitor experiments and analysis. CL generated most vectors and constructs. JR assisted with generation of CRISPR/Cas9 generated strains. WA and MR assisted with aldicarb behavioral assay. DO assisted with EMS screen and isolation of *dve-1* mutant. CB and MD aided in CloudMap bioinformatic analysis of the *uf171* mutant. MMF and KDA designed and interpreted results of all experiments and wrote the manuscript.

## Introduction

The mature human brain is composed of billions of neurons that are organized into functional circuits based on stereotyped patterns of synaptic connections that optimize circuit performance. Mature circuit connectivity is choreographed through a remarkable period of developmental circuit rewiring that is broadly conserved across species [1–4]. During this rewiring or remodeling phase, the mature circuitry is established through a tightly controlled balance: on the one hand, degenerative processes promote the elimination of juvenile synapses, while, on the other hand, maintenance or growth processes support the stabilization or formation of new connections. A combination of cell-intrinsic and extrinsic factors shape the progression of these events. For instance, activity-dependent microglial engulfment and elimination of synaptic material shapes connectivity of the retinogeniculate system in mice [5–8], while cell-intrinsic genetic programs such as the circadian clock genes Clock or Bmal1 influence GABAergic maturation and plasticity-related changes in the neocortex [9]. While molecular mechanisms supporting axon guidance and synapse formation have received considerable attention, our understanding of neuron-intrinsic molecular mechanisms controlling synapse elimination remains more limited. In particular, it is unclear how neuron-intrinsic synapse elimination processes are engaged in developing neural circuits. Improved mechanistic knowledge of these processes offers potential for important advances in our grasp of brain development. This knowledge may also inform the pathology underlying numerous neurodevelopmental diseases associated with altered connectivity and neurodegenerative diseases where synapse loss is a hallmark feature [10–13]. Indeed, recent work has suggested intriguing parallels between the elimination of synapses during development and neurodegenerative processes during disease [10–13].

The nematode *Caenorhabditis elegans* offers significant assets for addressing mechanistic questions about developmental neural circuit remodeling, particularly synapse elimination. *C. elegans* progresses through a highly stereotyped period of nervous system remodeling that establishes the neural connections characteristic of mature animals. 80 of the 302 neurons composing the adult nervous system, including 52 motor neurons, are born post-embryonically and integrated into pre-existing juvenile circuits following the first larval (L1) stage of development [14, 15]. The incorporation of these post-embryonic born motor neurons is accomplished following the first larval stage (L1) of development through a remarkable reorganization of circuit connectivity. One of the most striking aspects of this reorganization is the remodeling of synaptic connections in the GABAergic dorsal D-class (DD) motor neurons (**Figure 1A**) [16, 17]. Immediately after hatch, juvenile cholinergic synaptic inputs onto GABAergic DD neurons are located dorsally, while juvenile DD synaptic outputs onto muscles are located ventrally. During remodeling, the juvenile dorsal cholinergic synaptic inputs onto DD neurons are eliminated, and new synaptic inputs from post-embryonic-born presynaptic cholinergic neurons are established ventrally. In parallel, ventral DD GABAergic synaptic terminals are relocated dorsally, forming new GABAergic synaptic contacts onto dorsal muscles [18–22]. Though we now have a growing appreciation of the cellular processes that direct the post-embryonic redistribution of DD GABAergic outputs onto dorsal muscles, we have a limited understanding of how cholinergic inputs to DD neurons are remodeled. Prior work suggested a mechanism for antagonizing the remodeling of cholinergic inputs onto DD neurons through temporally controlled expression of the Ig domain family member OIG-1 [23, 24]; however, the mechanisms that promote remodeling of these inputs, in particular their elimination, have remained uncharacterized.

**Figure 1.**
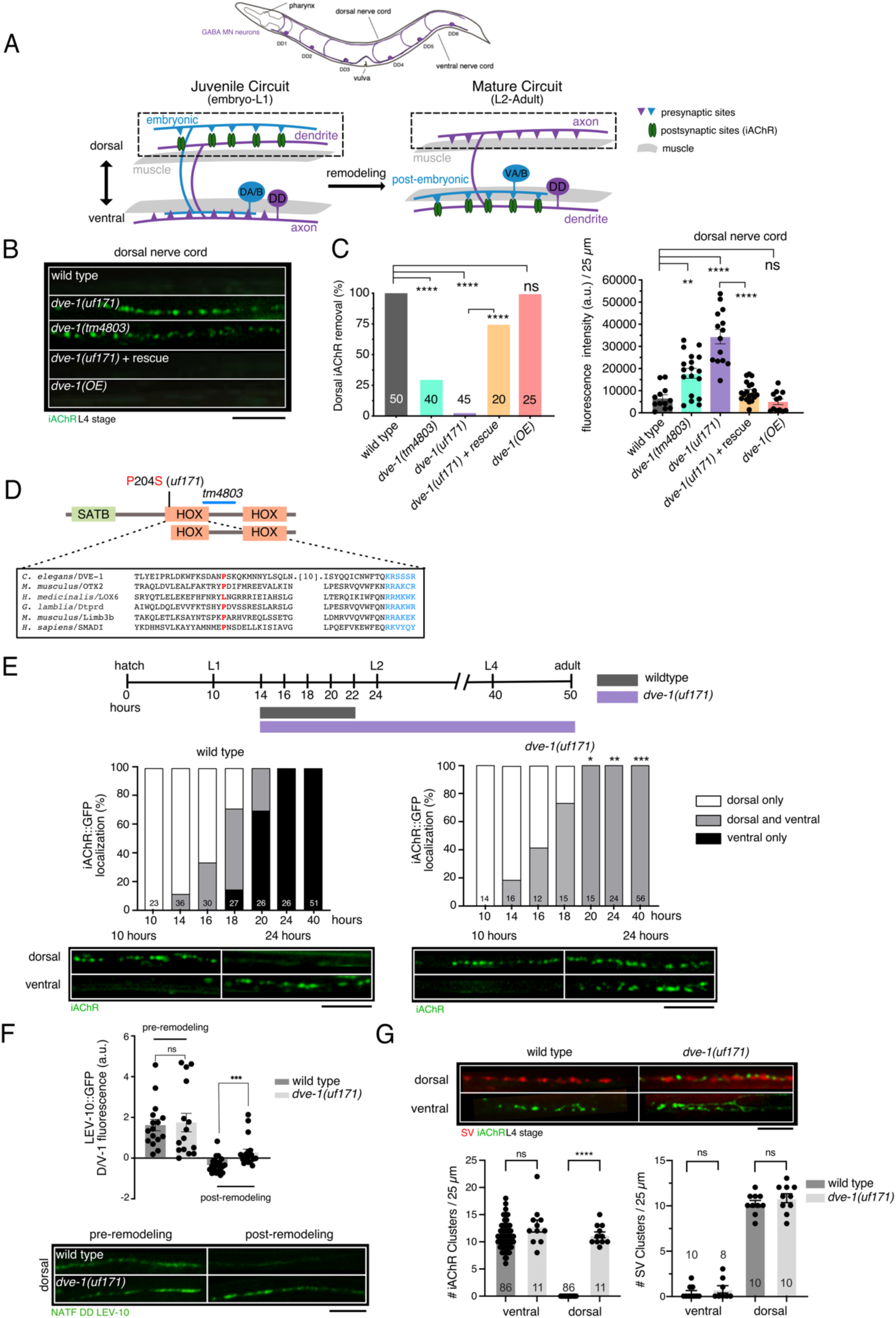
Mutations in the homeodomain transcription factor DVE-1 disrupt the removal of postsynaptic sites in GABAergic motor neurons. (A) Top, schematic of *C. elegans* labeling DD GABAergic motor neurons (purple). Bottom, schematic depicting motor circuit remodeling. Left, embryonic DD motor neurons (purple) receive input from cholinergic motor neurons (blue) in the dorsal nerve cord and make synaptic contacts onto ventral muscles. Right, after the L1-L2 molt, DD motor neurons are remodeled to receive inputs from postembryonic born cholinergic motor neurons in the ventral nerve cord and make synaptic contacts onto dorsal muscles. (B) Fluorescent confocal images of synaptic ACR-12::GFP clusters (iAChRs, green) in GABAergic DD processes of the dorsal nerve cord at L4 stage. By L4 stage, iAChR clusters are completely eliminated from the dorsal nerve cord of wild type animals. In contrast, iAChR clusters remain present in the dorsal nerve cord of *dve-1* mutants. Rescue refers to expression of genomic sequence encoding wild type DVE-1. OE refers to overexpression of *dve-1* in the wild type. In this and subsequent figures, iAChR refers to ACR-12::GFP unless otherwise indicated. Scale bar, 5 µm. (C) Quantification of iAChR clustering at L4 stage for the genotypes indicated. Left, bars represent the percentage of L4 stage animals where dorsal iAChRs have been completely removed. \*\*\*\**p*<0.0001, Fischer’s exact test with Bonferroni Correction. For this and subsequent figures, numbers within bars indicate the number of animals scored for each condition. Right, measures of average fluorescence intensity per 25 µm in the dorsal nerve cord. Bars indicate mean ± SEM. \*\**p*<0.05, \*\*\*\**p*<0.0001, ns – not significant ANOVA with Dunnett’s multiple comparisons test. Each point represents a single animal. The number of animals >10 per genotype. (D) Domain structure of DVE-1. SATB-like domain and homeodomains (HOX) are indicated. Site of substitution produced by *uf171* missense mutation (red) and region of *tm4803* deletion mutation (blue) are indicated. Inset, sequence alignment of homeodomain 1 (determined using NCBI Blast Conserved Domains). (E) Top, timeline of remodeling, approximate timing of transitions between larval stages and adulthood are indicated. Bars indicate duration of DD synaptic remodeling for wild type (blue) and *dve-1* mutants (magenta). Elimination of dorsal cord iAChR clusters is completed by 22 hours after hatch for wild type whereas dorsal iAChR clusters persist through adulthood in *dve-1* mutants. Middle, quantification of iAChR remodeling in DD neurons of wild type (left) and *dve-1* mutants (right). X-axis time from hatching in hours. Animals are binned according to the distribution of iAChR puncta as dorsal only (white), ventral only (black), or dorsal and ventral (grey). Bottom, representative images of dorsal and ventral iAChR clusters for wild type (left) and *dve-1* mutants (right) at the times indicated. \**p*<0.05, ** *p*<0.01, *** *p*<0.001, Fischer’s exact test. Scale bar, 5 µm. (F) Top, scatterplot of LEV-10::GFP dorsal/ventral fluorescence intensity ratio measurements per corresponding 25 µm regions of dorsal and ventral nerve cord expressed as dorsal/ventral fluorescence ratio -1. Values greater than zero indicate predominant dorsal localization while values less than zero indicate predominant ventral localization. The postsynaptic scaffold LEV-10::GFP is incompletely removed from dorsal nerve cord in *dve- 1(uf171)* mutants. Bars indicate mean ± SEM. \*\*\**p*<0.001, ns – not significant student’s t- test. Bottom, confocal images of LEV-10::GFP from the dorsal nerve cord before (pre- remodeling) and after (post-remodeling) the L1/L2 transition. Scale bar, 5 µm. NATF DD LEV-10 indicates tissue-specific labeling of endogenous LEV-10 [30]. Each point represents a single animal. The number of animals >10 per genotype. (G) Top, confocal images of the dorsal and ventral process from L4 stage wild type and *dve- 1(uf171)* mutants co-expressing the synaptic vesicle marker mCherry::RAB-3 (SV) and ACR-12::GFP marker (iAChR) in DD neurons. Scale bar, 5 µm. Bottom, quantification of iAChR clusters (left) and SV puncta (right) in dorsal and ventral processes of L4 stage DD neurons for wild type and *dve-1(uf171)* mutants. iAChRs in DD neurons are not eliminated from the dorsal nerve cord of L4 stage *dve-1* mutants, but new clusters form normally in the ventral nerve cord. SV clusters are properly reoriented from the ventral cord to the dorsal nerve cord in DD neurons of *dve-1* mutants. Bars indicate mean ± SEM. Student’s t-test, *\*\*\*\*p*<0.0001, ns – not significant. Each point represents a single animal.

We report the identification of a mechanism for neuron-intrinsic transcriptional control of synapse elimination during remodeling of the *C. elegans* motor circuit. From a forward genetic screen to isolate mutants whose juvenile postsynaptic sites on GABAergic DD neurons persist in mature animals, we obtained a mutation in the homeodomain transcriptional regulator *dve-1* that shares homology with mammalian special AT-rich sequence-binding (SATB) family members. We show that DVE-1 acts cell autonomously in GABAergic DD neurons to promote the removal of juvenile cholinergic synaptic inputs. Juvenile synaptic inputs are maintained into adulthood in *dve-1* mutants, leading to an accumulation of presynaptic cholinergic vesicles as well as alterations in circuit function and movement. We further show that precocious synapse elimination in *oig-1* mutants is reversed by mutation of *dve-1*, suggesting that DVE-1 promotes pro-degenerative processes that are antagonized by OIG-1. Our results reveal a novel neuron-intrinsic mechanism for developmentally timed synapse elimination through converging pro-degenerative and maintenance pathways.

## Results

### Distinct mechanisms direct developmental remodeling of presynaptic terminals versus postsynaptic sites in GABAergic neurons

Previous work by our lab and others showed that clusters of postsynaptic ionotropic acetylcholine receptors (iAChR) denote postsynaptic sites on DD neurons and undergo dorsoventral remodeling during the transition between the 1^st^ and 2^nd^ larval stages of development (L1/L2 transition) [23–25]. During this period, dorsal postsynaptic sites on DD neurons are removed, and the growth of new ventral postsynaptic sites is indicated by the appearance of newly formed ventral iAChR clusters. In animals co-expressing the synaptic vesicle marker mCherry::RAB-3 and the iAChR marker ACR-12::GFP, we found that the remodeling of cholinergic postsynaptic sites and GABAergic presynaptic terminals in DD neurons occurred with similar time courses, consistent with prior work (**Figure S1.1A,B**). Surprisingly, however, we found that mutations in several genes previously implicated in the remodeling of GABAergic presynaptic terminals had no appreciable effect on the remodeling of cholinergic postsynaptic sites in DD neurons (**Figure S1.1C, Table 1**). For example, juvenile mCherry::RAB-3 clusters were not fully removed from the ventral nerve cord of *ced-3/*caspase mutants at the L4 stage [26], indicating a failure to properly relocate GABAergic presynaptic terminals during remodeling. However, juvenile cholinergic postsynaptic sites were properly removed in DD neurons of *ced-3* mutants, as indicated by the absence of iAChR clusters in the dorsal nerve cord after the L2 stage (**Figure S1.1C**). Indeed, lingering synaptic vesicle clusters in the ventral nerve cord of *ced-3* mutants were interleaved with newly formed ventral iAChR clusters, demonstrating that the formation of new ventral postsynaptic sites during remodeling also occurs independently of *ced-3*. Mutations in several genes important for neurotransmitter release and calcium signaling also did not appreciably alter the remodeling of postsynaptic sites in DD neurons, though we noted clear delays in the remodeling of DD presynaptic terminals as found previously [19, 21] (**Table 2**). Of the genes we tested, only mutation of the RyR/*unc-68* gene produced a modest delay in the remodeling of postsynaptic sites, suggesting calcium release from intracellular stores contributes (**Table 2**). Taken together, our findings demonstrate that mechanisms for remodeling postsynaptic sites in DD neurons are distinct from those previously implicated in the remodeling of DD GABAergic presynaptic terminals.

**Table 1.**
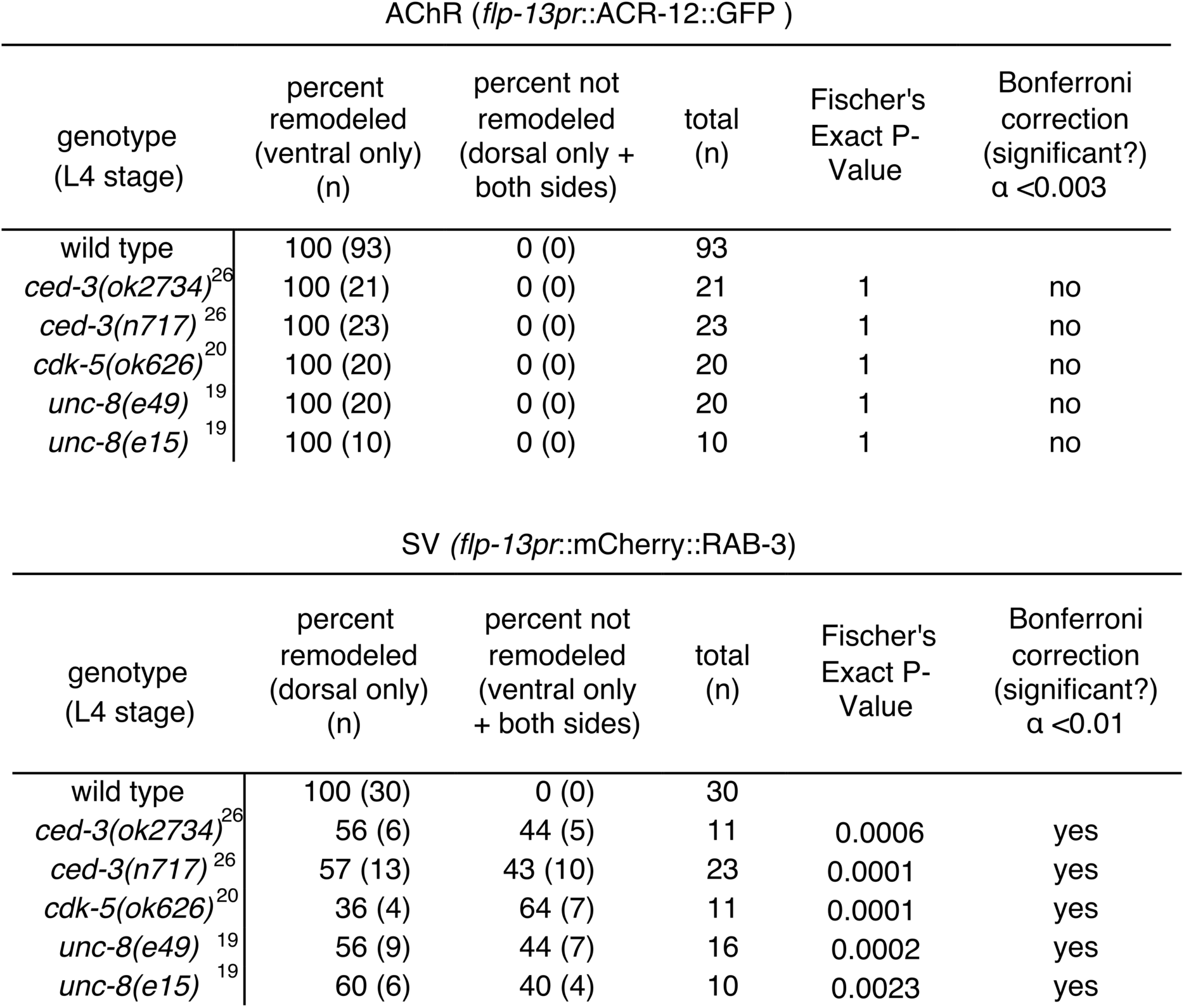
Mutations that delay GABAergic presynaptic remodeling do not affect cholinergic postsynaptic remodeling in DD motor neurons

**Table 2.** Effects of synaptic activity and calcium signaling on synaptic remodeling

| AChR (*flp-13pr::ACR-12::GFP*) |  |  |  |  |  |
| --- | --- | --- | --- | --- | --- |
| genotype<br>(24hrs post-hatch) | percent<br>remodeled<br>(ventral only)<br>(n) | percent not<br>remodeled<br>(dorsal only +<br>both sides) | total<br>(n) | Fischer's<br>Exact P-Value | Bonferroni<br>correction<br>(significant?)<br>$\alpha < 0.0055$ |
| wild type | 96 (64) | 4 (3) | 67 |  |  |
| *unc-2(e55)*<sup>19</sup> | 100 (10) | 0 (0) | 10 | 1 | no |
| *cca-1(ad1650)* | 100 (17) | 0 (0) | 17 | 1 | no |
| *itr-1(sa73)* | 100 (18) | 0 (0) | 18 | 1 | no |
| *unc-68(e540)* | 50 (6) | 50 (6) | 12 | 0.0002 | yes |
| *unc-13(e51)* | 100 (11) | 0 (0) | 11 | 1 | no |
| *unc-43(e408)* | 95 (19) | 5 (1) | 20 | 1 | no |
| *unc-18(e234)*<sup>21</sup> | 100 (10) | 0 (0) | 10 | 1 | no |
| *unc-17(e113)* | 76 (13) | 24 (4) | 17 | 0.0284 | no |

| genotype<br>(L4 stage) | percent<br>remodeled<br>(ventral only)<br>(n) | percent not<br>remodeled<br>(dorsal only +<br>both sides) | total<br>(n) | Fischer's<br>Exact P-Value | Bonferroni<br>correction<br>(significant?)<br>$\alpha < 0.003$ |
| --- | --- | --- | --- | --- | --- |
| wild type | 100 (93) | 0 (0) | 93 |  |  |
| *unc-2(e55)*<sup>19</sup> | 100 (10) | 0 (0) | 10 | 1 | no |
| *unc-2(zf35)*<sup>19</sup> | 86 (6) | 14 (1) | 7 | 0.07 | no |
| *cca-1(ad1650)* | 100 (10) | 0 (0) | 10 | 1 | no |
| *itr-1(sa73)* | 100 (10) | 0 (0) | 10 | 1 | no |
| *unc-68(e540)* | 100 (11) | 0 (0) | 11 | 1 | no |
| *unc-43(e408)* | 100 (9) | 0 (0) | 9 | 1 | no |
| *unc-43(n498)* | 100 (14) | 0 (0) | 14 | 1 | no |
| *unc-18(e234)*<sup>21</sup> | 100 (8) | 0 (0) | 8 | 1 | no |
| *unc-31(e928)*<sup>21</sup> | 100 (10) | 0 (0) | 10 | 1 | no |
| *unc-17(e113)* | 100 (11) | 0 (0) | 11 | 1 | no |

| SV (*flp-13pr::mCherry::RAB-3*) |  |  |  |  |  |
| --- | --- | --- | --- | --- | --- |
| genotype<br>(24hrs post-hatch) | Percent<br>remodeled<br>(dorsal only)<br>(n) | percent not<br>remodeled<br>(ventral only<br>+ both sides) | total<br>(n) | Fischer's<br>Exact P-Value | Bonferroni<br>correction<br>(significant?)<br>$\alpha < 0.025$ |
| wild type | 97 (28) | 3 (1) | 29 |  |  |
| *unc-18(e234)*<sup>21</sup> | 40 (4) | 60 (6) | 10 | 0.0004 | yes |
| *unc-17(e113)* | 33 (2) | 67 (4) | 6 | 0.0014 | yes |

### Identification of *dve-1* as a novel transcriptional regulator of synapse elimination

Motivated by these findings, we pursued a forward mutagenesis screen to identify novel mechanisms controlling the remodeling of postsynaptic sites in DD neurons. Specifically, we screened to obtain mutants where juvenile postsynaptic sites, indicated by dorsal iAChR clusters, were not properly eliminated during remodeling and instead persisted into maturity (**Figure S1.2A,B**), and isolated a recessive mutant, *uf171*. Dorsal postsynaptic sites are normally eliminated before the L2 stage (22 hours after hatch) in wild type but remained visible through the late L4 stage (>40 hours after hatch) in *uf171* mutants (**Figure 1B,C,E; Figure S1.3A**). Whole genome sequence analysis of *uf171* mutants revealed a point mutation in the gene encoding the homeodomain protein DVE-1, producing a proline to serine (P/S) substitution (**Figure 1D**). Expression of the wild type *dve-1* gene in *dve-1(uf171)* mutants restored normal elimination of juvenile postsynaptic sites (**Figure 1B,C; Figure S1.3A),** while *dve-1* overexpression in wild type animals did not produce appreciable changes in removal (**Figure 1B,C; Figure S1.3A**). A similar failure in the elimination of juvenile postsynaptic sites was also evident in another available *dve- 1* mutant, *dve-1(tm4803)*, that harbors a small insertion/deletion mutation (**Figure 1B,C; Figure S1.3A**). DVE-1 was of particular interest as it is a homeodomain transcription factor sharing homology with mammalian SATB transcription factors and was previously implicated in the regulation of a mitochondrial stress response in *C. elegans* [27]. The P/S substitution encoded by *dve-1(uf171)* affects a highly conserved proline residue predicted to lie within a loop between helices I and II of the first homeobox domain of *dve-1*. *dve-1(tm4803)* deletes a portion of predicted helix III in the same homeobox domain and a splice site leading to 65 bp insertion (**Figure 1D**) [28]. As *dve-1* null mutants are embryonic lethal [27], both mutations are predicted to be hypomorphic.

The postsynaptic scaffold protein LEV-10 is associated with cholinergic postsynaptic sites in body wall muscles and GABAergic neurons [29, 30]. Using a previously developed strategy for cell-specific labeling of endogenous LEV-10 [30], we found that mutation of *dve-1* also slowed the removal of LEV-10 from the dorsal processes of DD neurons during remodeling. LEV-10 was primarily associated with dorsal processes prior to remodeling and was redistributed to ventral processes during remodeling (**Figure 1F**). Similar to juvenile iAChR clusters, dorsal LEV-10 scaffolds in GABAergic neurons of *dve-1* mutants were not properly eliminated during remodeling, further demonstrating that DVE-1 coordinates the removal of juvenile postsynaptic sites in DD neurons. Interestingly, mutation of *dve-1* did not significantly affect the formation of new ventral postsynaptic sites on DD neurons during remodeling or their stability (**Figure 1E,G; Figure S1.3B**). Likewise, mutation of *dve-1* had little effect on the density of recently characterized DD dendritic spines that are formed after remodeling (**Figure S1.3C,D**) [31–33]. These findings indicate that the evolutionarily conserved transcription factor DVE-1 specifically governs the elimination of juvenile postsynaptic sites during remodeling without affecting the formation or maturation of new postsynaptic sites. Notably, the remodeling of DD GABAergic presynaptic terminals also occurred normally in *dve-1* mutants (**Figure 1G**), further indicating that distinct neuron-intrinsic programs direct remodeling of the pre- and postsynaptic domains of GABAergic DD neurons (**Figure 1G; Figure S1.1C; Tables 1, 2**).

### Lingering iAChRs in *dve-1* mutants are organized into structural synapses

To test if lingering juvenile postsynaptic sites in the dorsal nerve cord of *dve-1* mutants are organized into structurally intact synapses, we first asked whether these iAChR clusters were localized at the cell surface. Using an established approach for *in vivo* antibody labeling of cell surface receptors [33, 34], we found that lingering iAChR clusters were visible in the dorsal cord of *dve-1* mutants, but not in controls (**Figure 2A**). We also found that most iAChR clusters retained in dorsal GABAergic DD processes of *dve-1* mutants were closely apposed by synaptic vesicle assemblies of cholinergic DA/B axons in the dorsal nerve cord (**Figure 2B**), suggesting incorporation into structural synapses.

**Figure 2.**
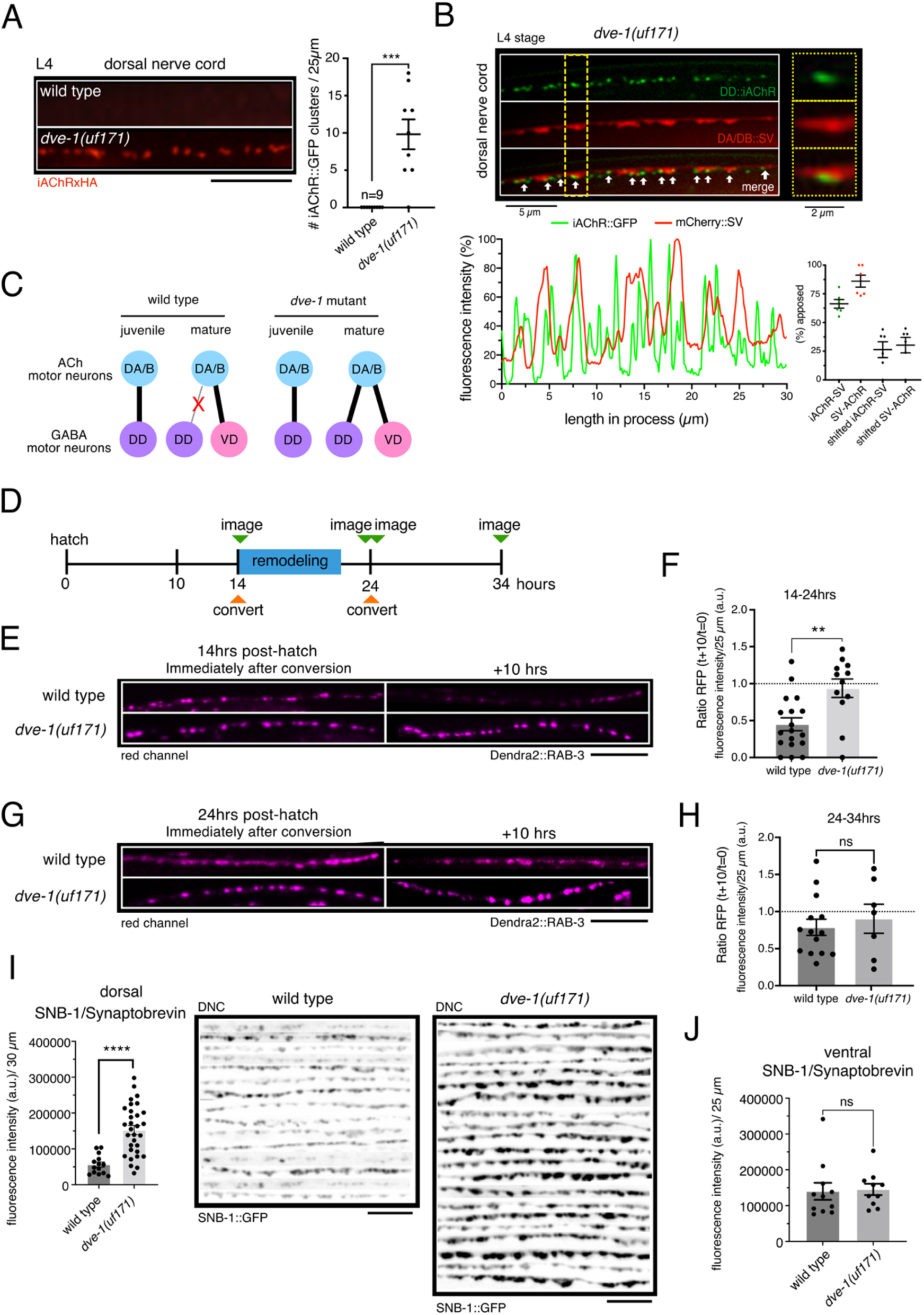
Mutation of *dve-1* enhances stability of cholinergic presynaptic sites. (A) Left, fluorescent confocal images of L4 stage cell surface iAChR clusters in the dorsal nerve cord labeled by anti-HA antibody fluorescence (red). Scale bar, 5 µm. Right, scatterplot of average number of dorsal receptor clusters per 25 µm. Each point represents a single animal n=9 for both genotypes. Data points represent mean ± SEM. \*\*\**p*<0.001, students t- test. (B) Top, merged fluorescent confocal images of L4 stage *dve-1* mutant dorsal nerve cord showing lingering clusters of juvenile postsynaptic iAChRs (*flp-13pr*::ACR-12::GFP, green) in the DD neurons and synaptic vesicles (SV, *unc-129pr*::mCherry::RAB-3, red) in presynaptic cholinergic DA/DB neurons. Dashed yellow line indicates area shown in inset (right). Bottom, line scan (left) and quantification (right) indicating % apposition of postsynaptic iAChR and presynaptic cholinergic vesicle clusters. Line scan indicates relative fluorescence intensity of iAChR (green) and SV (red) for a 30 µm region of L4 stage *dve-1* mutant dorsal nerve cord. Scatterplot shows the percentage apposition between iAChR clusters in DD motor neurons and cholinergic synaptic vesicle puncta in DA/B motor neurons (green). As controls (black) each iAChR and SV line scan was shifted by 2 µm and reassessed to determine percent apposed by chance. Each point represents a single animal. The number of animals >5. Data points indicate mean ± SEM. (C) Simplified schematic of DA/B synaptic output. In the juvenile circuit (L1 stage) DA/B motor neurons (blue) synapse onto DD motor neurons (purple) in both wild type (left) and *dve-1* mutants (right). In the wild type mature circuit (L2 through adulthood) DA/B connections with DD motor neurons have been removed, and new connections with postembryonic born VD motor neurons (pink) are formed (left). Grey line with red “X” indicates removal of juvenile connection. Our findings suggest that connections between cholinergic DA/B neurons and DD neurons are retained in *dve-1* mutants retain (right). (D) Schematic of photoconversion experimental design. Animals were imaged 14 hours after hatch, prior to and immediately after photoconversion. Photoconverted animals were imaged again 10 hours later. Photoconversion was performed again 24 hours after hatch. Images were captured immediately after conversion and again 10 hours later (34 hours after hatch). (E) Confocal images of the dorsal nerve cord of wild type (top) and *dve-1(uf171)* mutants (bottom) showing red Dendra2::RAB-3 clusters in cholinergic DA/B neurons (*unc- 129pr*::Dendra2::RAB-3) either immediately after photoconversion from green to red at 14 hours after hatch (left) or 10 hours later (right). Juvenile Dendra2::RAB-3 clusters are largely removed during wild type remodeling but are more stable in *dve-1* mutants. Scale bar, 5 µm. (F) Scatterplot showing red Dendra2::RAB-3 fluorescence intensity 10 hours after photoconversion normalized to fluorescence intensity immediately after photoconversion prior to remodeling for wild type (left) and *dve-1(uf171)* mutants (right). Each point indicates a single animal. The number of animals >10 per genotype. Bars indicate mean ± SEM. *\*\*p*<0.01, student’s t-test. (G) Confocal images of the dorsal nerve cord of wild type (top) and *dve-1(uf171)* mutants (bottom) showing red Dendra2::RAB-3 clusters in cholinergic DA/B neurons either immediately after photoconversion from green to red at 24 hours after hatch (left) or 10 hours later (right). For both wild type and *dve-1* mutants, photoconverted cholinergic Dendra2::RAB-3 clusters are more stable following remodeling compared with during remodeling. Scale bar, 5 µm. (H) Scatterplot showing red Dendra2::RAB-3 fluorescence intensity 10 hours following photoconversion normalized to fluorescence intensity immediately after photoconversion at 24 hours after hatch (after remodeling) for wild type (left) and *dve-1(uf171)* mutants (right). Each point indicates a single animal. The number of animals >6 per genotype. Bars indicate mean ± SEM. ns - not significant, student’s t-test. (I) Left, scatterplot showing average SNB-1::GFP fluorescence intensity in L4 stage cholinergic neurons (*acr-5pr*::SNB-1::GFP) of the dorsal nerve cord (DNC) for wild type and *dve-1* mutants. Each point indicates a single animal. The number of animals >10 per genotype. Bars indicate mean ± SEM. \*\*\*\**p*< 0.0001, student’s t-test. Right, stacked fluorescent images of the dorsal nerve cord showing SNB-1::GFP clusters in cholinergic neurons of L4 stage wild type and *dve-1(uf171)* mutants. Images on each line are from different animals. Scale bar, 5 µm. (J) Scatterplot of average synaptic vesicle puncta (SNB-1::GFP) per 30 µm in the ventral nerve cord of VB motor neurons of L4 stage wild type and *dve-1* mutant animals. Each point represents a single animal. At least 10 animals per genotype. Bars indicate mean ± SEM. ns - not significant, student’s t-test.

During remodeling, cholinergic DA/B connections with DD neurons in the dorsal nerve cord are removed and new DA/B connections are established with post-embryonic born ventrally directed GABAergic D-class (VD) motor neurons (**Figure 1A**, **Figure 2C**). To investigate how the remodeling of presynapses within the cholinergic DA/B axons may be affected by mutation of *dve- 1*, we expressed the photoconvertible synaptic vesicle marker Dendra2::RAB-3 in cholinergic DA/B neurons (**Figure 2D**). We first examined the distribution of Dendra2::RAB-3 in the wild type dorsal nerve cord immediately prior to the onset of DD remodeling (approximately 14 hours after hatch) (**Figure 2E**). Prior to photoconversion, clusters of green Dendra2::RAB-3 fluorescence were distributed along the length of cholinergic axons in the dorsal nerve cord. Brief exposure of the dorsal nerve cord to 405 nm light produced immediate and irreversible photoconversion of Dendra2::RAB-3 signals from green to red fluorescence (**Figure S2.1E**). In wild type, Dendra2::RAB-3 clusters that had been photoconverted to red fluorescence prior to the onset of remodeling were strikingly reduced following remodeling (10 hours later, 55 ± 9% reduction) and were replaced by new synaptic vesicle clusters (green fluorescence) (**Figure 2E,F; Figure S2.1A,B,E**). In contrast, wild type Dendra2::RAB-3 clusters photoconverted after completion of remodeling (at approximately 24 hours after hatch) remained largely stable over the subsequent 10 hours (**Figure 2G-H**, **Figure S2.1F**). We also noted the appearance of green RAB-3 clusters during this time frame (24-34 hours after hatch), indicating parallel addition of new vesicular material (**Figure S2.1C,D,F**). Thus, synaptic vesicle clusters in wild type DA/B axons are largely removed and replaced during the 10-hour period of remodeling but are more stable over a 10- hour time window immediately following completion of remodeling, offering intriguing evidence for developmental stage-specific regulation of cholinergic synaptic vesicle stability. Importantly, this transient reorganization of presynaptic release sites in wild type cholinergic axons occurs concurrently with the removal and remodeling of postsynaptic sites in DD GABAergic neurons.

We noted a striking change in the stability of synaptic vesicle material in cholinergic axons of *dve- 1* mutants during remodeling. Dorsal cholinergic Dendra2::RAB-3 clusters photoconverted prior to the onset of synaptic remodeling were preserved throughout remodeling in *dve-1* mutants (**Figure 2E,F; Figure S2.1E**), indicating enhanced stability of cholinergic terminals presynaptic to DD neurons. The addition of new synaptic vesicles during this time frame (14-24 hours after hatch) was not appreciably affected by mutation of *dve-1*, as indicated by similar increases in green Dendra2::RAB-3 fluorescence across wild type and *dve-1* mutant cholinergic axons (**Figure S2.1A,B,E**). RAB-3 clusters that were photoconverted after the completion of remodeling (approximately 24 hours after hatch) remained detectable 10 hours later in *dve-1* mutants, also similar to wild type (**Figure 2G,H; Figure S2.1F**). Dorsal axon green Dendra2::RAB-3 fluorescence was increased by roughly 2-fold in *dve-1* mutants compared to wild type 24-34 hours after hatch, suggesting enhanced addition or stabilization of new synaptic vesicles at *dve-1* mutant cholinergic axon terminals after remodeling (**Figure S2.1C,D,F**). To explore this in more detail, we examined the distribution of the synaptic vesicle marker SNB-1::GFP in dorsal cholinergic axons of L4 stage *dve-1* mutants. The intensity of SNB-1::GFP clusters was increased in dorsal cholinergic axons of *dve-1* mutants compared to wild type (**Figure 2I**), whereas SNB-1::GFP fluorescence intensity in ventral cholinergic axons of *dve-1* mutants was unchanged (**Figure 2J**). These data suggest DVE-1 specifically regulates cholinergic synaptic contacts onto DD neurons (**Figure S2.1E**). Notably, UNC-10::GFP fluorescence intensity (labeling active zones) in L4 stage dorsal cholinergic axons was not appreciably affected by *dve-1* mutation (**Figure S2.1E**). Thus, mutation of *dve-1* leads to an increase in the stability or recruitment of synaptic vesicle material at dorsal cholinergic axon terminals but does not appreciably alter the size or density of active zones. Together with our previous findings, these data suggest that wild type *dve-1* promotes destabilization of both vesicle assemblies in presynaptic cholinergic axons and cholinergic postsynaptic sites in GABAergic neurons during remodeling.

### A failure to eliminate postsynaptic sites via DVE-1 leads to enhanced activity and altered motor behavior

To investigate how a failure of synapse elimination may impact circuit function, we first sought to determine whether preserved structural connections between dorsal cholinergic axons and GABAergic DD neurons of *dve-1* mutants were functional in adults. To address this question, we used combined cell-specific expression of Chrimson for cholinergic depolarization [35, 36] and GCaMP6 for monitoring [Ca^2+^] changes in the postsynaptic GABAergic motor neurons [37] (**Figure S3.1A**). We recorded Ca^2+^ transients from young adult GABAergic DD or VD motor neurons in response to presynaptic DA/B cholinergic depolarization. We did not observe an appreciable change in the magnitude of stimulus-elicited Ca^2+^ transients in VD neurons but noted striking changes in DD neurons. Specifically, we found that photostimulation elicited a modest Ca^2+^ response in roughly 37% of wild type DD neurons tested, consistent with a low degree of synaptic connectivity between these neurons in adults as predicted by the wiring diagram [15, 38]. The percentage of responsive DD neurons (85%) and the average magnitude of stimulus-elicited Ca^2+^ transients increased significantly for *dve-1* mutants (**Figure 3A**), demonstrating enhanced functional connectivity between dorsal cholinergic neurons and GABAergic DD neurons of adult *dve-1* mutants.

**Figure 3.**
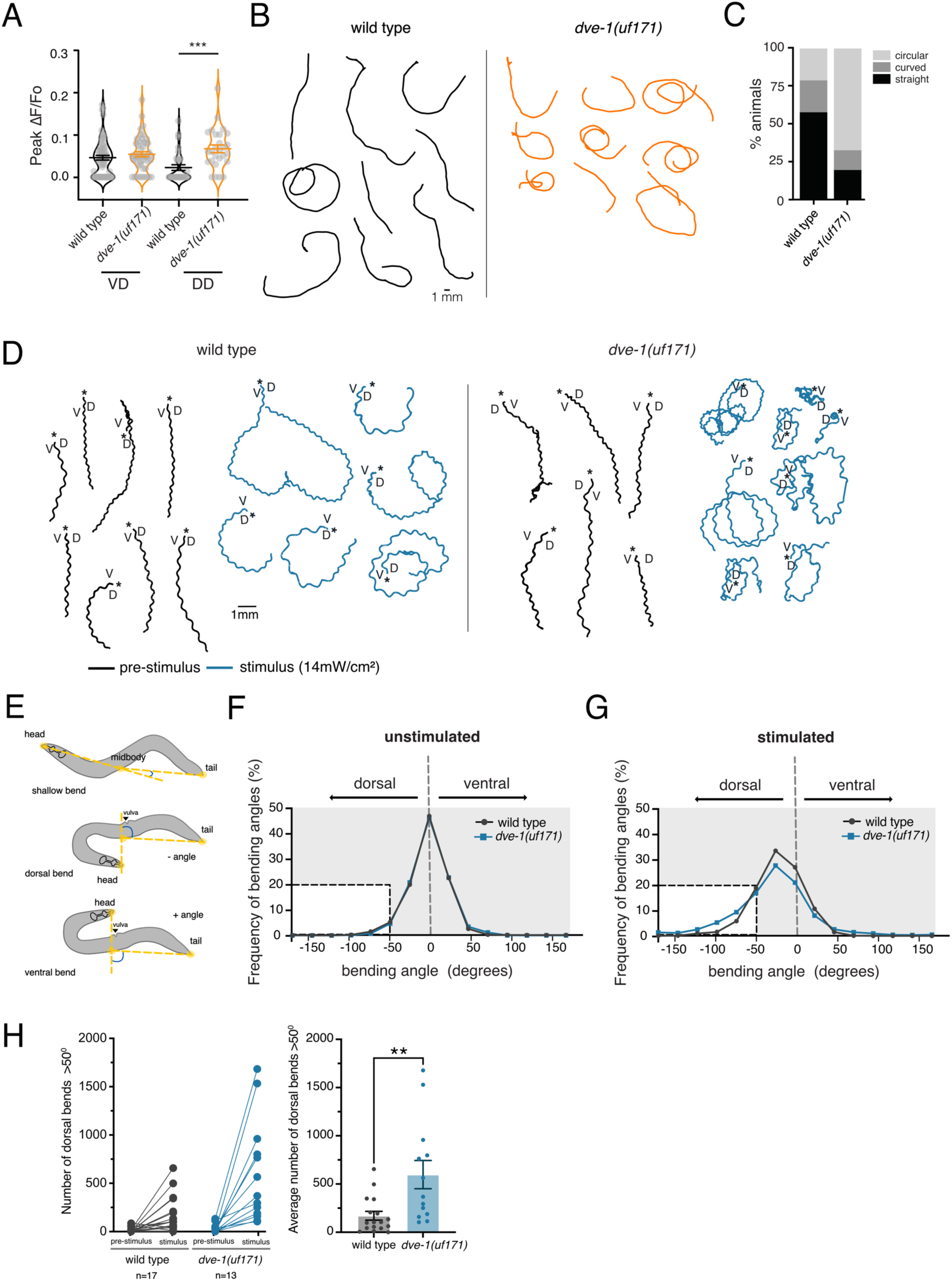
Failure of synapse elimination in *dve-1* mutants produces dorsal turning bias. (A) Scatter plot showing peak calcium response (ΔF/F_0_) in DD and VD GABAergic neurons to photostimulation of presynaptic DA and DB cholinergic neurons for wild type and *dve-1(uf171)* mutants. Horizontal bars indicate mean peak ΔF/F_0_ ± SEM. *** *p*<0.001, ANOVA with Tukey’s multiple comparison. n=16; animals for each condition. Number of cells quantified: wt DD: 30, *dve-1(uf171)* DD: 27, wt VD: 64, *dve-1(uf171)* VD: 47. (B) Representative locomotion tracks for wild type (black) and *dve-1(uf171)* (orange) animals recorded over 5 minutes of single worm tracking on NGM OP50 plates. Scale bar, 1 mm. (C) Percentage of straight, curved, or circling tracks for wild type and *dve-1(uf171)* mutants. (D) Tracks for wild type (left) and *dve-1* mutant (right) animals during forward runs (30 s) prior to or during photostimulation. Asterisks indicate start of track. D/V indicate dorsal and ventral directions. (E) Schematics of bending angle measurements. Solid orange circles indicate the vertices (head, midbody, and tail) of the body bending angle (blue) measured. (F) Frequency distribution of body bending angles measured prior to photosimulation for wild type (black) and *dve-1(uf171)* (blue). Negative bending angle values indicate dorsal, while positive bending angle values indicate ventral. Inset highlights bending events greater than 50°. wild type n=17, *dve-1(uf171)* n=13. (G) Frequency distribution of body bending angles measured during photostimulation for wild type (black) and *dve-1(uf171)* (blue). Negative bending angle values indicate dorsal, while positive bending angle values indicate ventral. Inset highlights bending events greater than 50°. wild type n=17, *dve-1(uf171)* n=13. (H) Left, scatterplot of total number of dorsal bends greater than 50° before and after photostimulation. Points with connecting line represents a single animal. Right, average number of dorsal bends greater than 50° during the period of photostimulation for wild type and *dve-1(uf171)* animals. Bars indicate mean ± SEM. \*\**p*<0.01, student’s t-test. wild type n=17, *dve-1(uf171)* n=13.

We next asked how altered functional connectivity in the motor circuit of *dve-1(uf171)* mutants might affect circuit performance and behavior. Automated tracking of single worms during exploratory behavior showed *dve-1(uf171)* mutants frequently moved in loose, dorsally directed circles, whereas wild type animals were more likely to adopt straight trajectories (**Figure 3B**). During 5 minutes of continuous tracking, roughly 80% of *dve-1(uf171)* mutants circled or curved, approximately 60% of these in the dorsal direction, while only 20% of wild type circled (**Figure 3B,C**). The dorsal circling behavior of *dve-1* mutants suggested that altered synaptic output from the motor circuit may produce a turning bias. In addition to GABAergic neurons, dorsally directed DA/B cholinergic motor neurons are presynaptic to body wall muscles. We speculated that the increased abundance of cholinergic synaptic vesicles in dorsal motor axons of *dve-1* mutants may enhance cholinergic activation of dorsal muscles and elicit more robust dorsal turning. In support of this idea, we found that *dve-1* mutants were hypersensitive to the paralyzing effects of the acetylcholinesterase inhibitor aldicarb, an indicator of elevated acetylcholine release [39] (**Figure S3.1B**).

To explore this further, we tracked animals during depolarization of dorsal cholinergic neurons by cell-specific photoactivation using Chrimson. Prior to stimulation, control animals moved in predominantly forward trajectories (**Figure 3D-F**). As expected, photostimulation of DA/B motor neurons (625 nm, 14 mW/cm^2^) enhanced dorsal turning in control animals, often leading to large dorsally oriented circles (**Figure 3D,G**). DA/B motor neuron photostimulation elicited heightened turning responses in *dve-1* mutants, increasing dorsal turns by ∼2.5 fold compared with photostimulation in wild type and leading to tight dorsally oriented circles (**Figure 3D-H**). The enhanced dorsal turning of *dve-1* mutants was associated with an increase in the depth of dorsal bends compared to wild type (**Figure 3G,H, Figure S3.1C,D**) and was not observed in animals that lacked Chrimson expression or in the absence of retinal chromophore (**Figure S3.1E,F**). Chrimson expression was also not appreciably different across *dve-1* mutants and controls (**Figure S3.1G**). We propose that increased acetylcholine release onto dorsal muscles of *dve-1* mutants enhances dorsal bending and circling, perhaps due to an increase in the size of the synaptic vesicle pool in dorsal cholinergic axons. This interpretation is consistent with the increased abundance of synaptic vesicle material in dorsally projecting cholinergic axons of *dve- 1* mutants. Ectopic activation of dorsally projecting GABAergic DD neurons in *dve-1* mutants might be expected to enhance dorsal inhibition, countering the effects of dorsal excitation. However, we speculate that the number and strength of synaptic connections from dorsal cholinergic motor neurons to dorsal body wall muscles overwhelms any increase in dorsal inhibition. Together, our results suggest mutation of *dve-1* impacts functional connectivity both through retention of juvenile connectivity onto DD motor neurons and through an increase in cholinergic transmission onto dorsal muscles.

### Synapse elimination occurs through a convergence of DVE-1 regulated destabilization and removal of OIG-1 antagonism

The timing of DD neuron remodeling is, in part, determined through temporally controlled expression of the Ig-domain protein OIG-1 [24, 33]. The expression of an *oig-1pr::*GFP transcriptional reporter in L1 stage DD neurons was not appreciably altered in *dve-1* mutants (**Figure 4A**), in alignment with prior evidence that *oig-1* expression is regulated independently of *dve-1*. The Ig domain protein OIG-1 normally antagonizes synaptic remodeling. In *oig-1* mutants, the remodeling of postsynaptic sites in DD neurons, including both elimination of dorsal juvenile postsynaptic sites and the formation of new ventral postsynaptic sites, occur precociously compared with wild type [23, 24]. However, unlike the precocious removal of dorsal postsynaptic sites in DD neurons of *oig-1* mutants, the removal of dorsal postsynaptic sites in *oig-1*;*dve-1* double mutants was strikingly delayed. The juvenile dorsal postsynaptic sites were retained in DD neurons of *oig-1*;*dve-1* mutants through late L4 stage, similar to *dve-1* single mutants (**Figure 4B-C,F**). DVE-1 is therefore required for synapse elimination even under conditions where antagonistic processes promoting synapse stabilization are disrupted. Conversely, similar to the premature formation of new ventral postsynaptic sites in *oig-1* single mutants, ventral synapses formed precociously in *oig-1*;*dve-1* double mutants (**Figure 4D-E, F**). Thus, disruption of *dve-1* function reversed precocious synapse elimination in *oig-1* mutants but did not impact the premature assembly of ventral postsynaptic sites, supporting the independence of programs for synapse elimination versus growth and suggesting independent functions for OIG-1 in each (**Figure 4F,G**). Overall, our findings show that mature connectivity is sculpted through a convergence of DVE-1 regulated elimination processes and temporally regulated OIG-1 based stabilization mechanisms.

**Figure 4.**
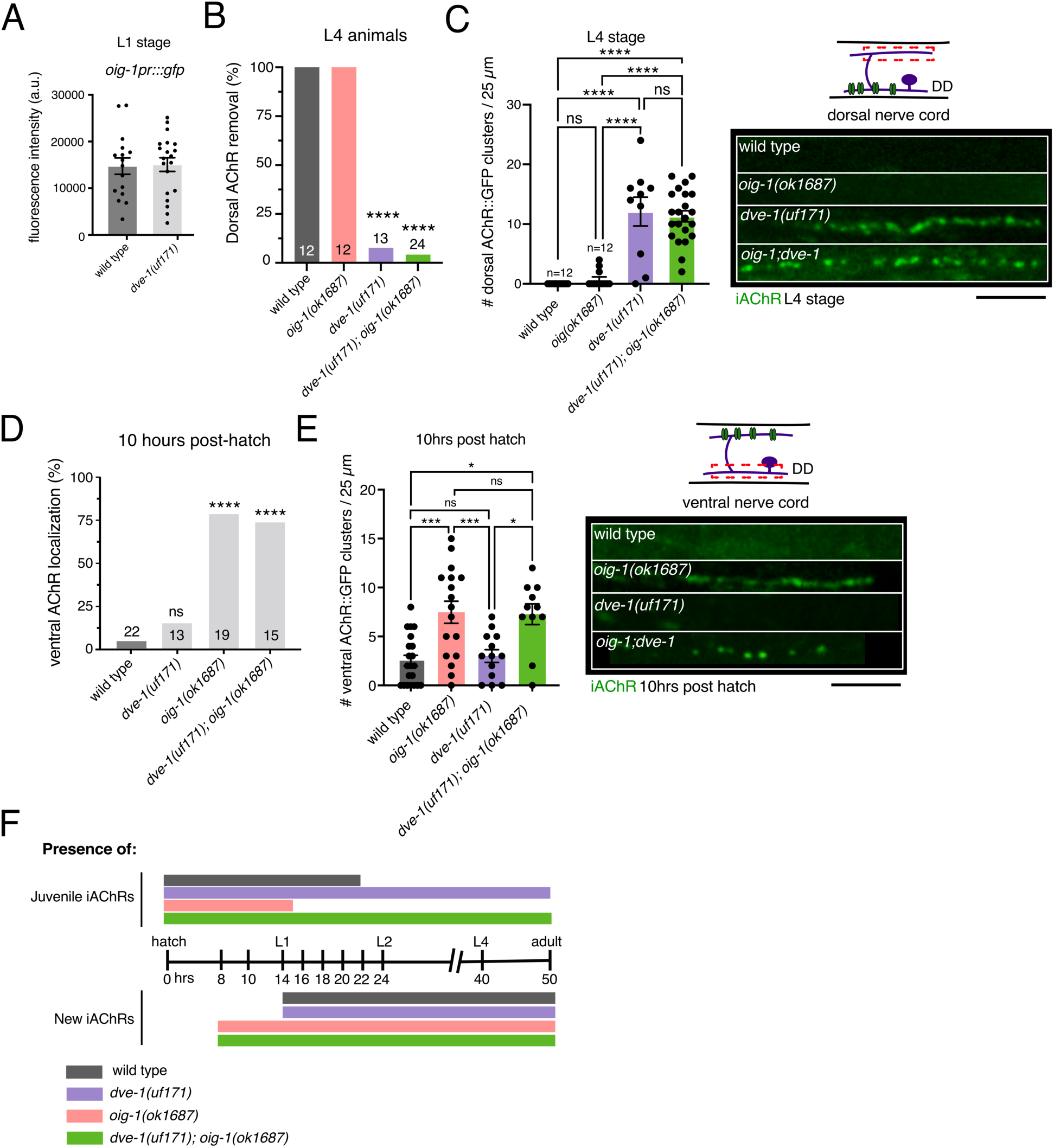
*dve-1* synaptic destabilization acts in parallel to *oig-1* antagonism of remodeling. (A) Average fluorescence intensity of *oig-1pr::gfp* in DD soma of L1 stage wild type and *dve- 1* mutants. Each point represents a single DD cell body. Imaged 2 DD neurons/animal. Wild type n=8, *dve-1* mutants n=10. Bars indicate mean ± SEM. (B) The percentage of animals where dorsal iAChRs are eliminated for L4 stage wild type, *oig-1(ok1687)*, *dve-1(uf171)* and *dve-1(uf171);oig-1(ok1687)* double mutants. \*\*\*\**p*< 0.0001, Fischer’s exact test with Bonferroni Correction. (C) Left, quantification of average number of iAChR clusters in L4 stage DD neurons per 25 µm of the dorsal nerve cord for the genotypes indicated. Each dot represents a single animal, at least 10 animals per genotype. Bars indicate mean ± SEM. \*\*\*\**p*<0.0001, ns - not significant, ANOVA with Dunnett’s multiple comparisons test. Right, fluorescent confocal images of iAChR clusters in L4 stage DD neurons of the dorsal nerve cord for the genotypes indicated. The precocious removal of dorsal iAChRs in *oig-1* mutants is reversed by mutation of *dve-1*. iAChR clusters persist through L4 stage in in both *dve-1* single and *oig-1;dve-1* double mutants. Scale bar, 5 µm. (D) Quantification of iAChR ventral localization in DD neurons in wild type, *oig-1(ok1687)* mutant, *dve-1(uf171)* mutant and *dve-1(uf171), oig-1(ok1687)* double mutant L1 stage animals (10 hours). \*\*\*\**p*<0.0001, ns-not significant, Fischer’s exact test. (E) Left, quantification of average number of iAChR clusters in DD neurons per 25 µm of the ventral nerve cord at L1 stage for the genotypes indicated. Each dot represents a single animal, at least 10 animals per genotype. Bars indicate mean ± SEM. ANOVA with Dunnett’s multiple comparisons test, \**p*<0.05, \*\*\**p*<0.001, ns-not significant. Right, fluorescent confocal images of iAChR clusters in the ventral processes of L1 stage DD neurons. Growth of ventral iAChR cluster occurs precociously in *oig-1* mutants and is unaffected by *dve-1* mutation. Scale bar, 5 µm. (F) Timeline of development, approximate timing of transitions between larval stages and to adulthood are indicated. Bars indicate the presence of juvenile dorsal iAChRs (top) or ventral iAChRs formed during remodeling (bottom). *dve-1* mutants (magenta) fail to remove juvenile synapses but form ventral synapses normally. *oig-1* mutants (orange) show precocious removal of juvenile synapses and precocious formation of ventral synapses compared to wild type (blue). *oig-1;dve-1* double mutants (green) fail to remove juvenile synapses similar to *dve-1* single mutants, but show precocious formation of new ventral synapses similar to *oig-1* single mutants.

### DVE-1 localization to GABAergic nuclei is required for synapse elimination

To better understand potential mechanisms for DVE-1 actions in synapse elimination, we examined *dve-1* expression using a strain where a GFP coding sequence was inserted in the *dve-1* genomic locus [40]. As noted previously [40], we observed prominent expression of endogenously labeled DVE-1::GFP in intestinal cells. However, we also noted neuronal expression of DVE-1::GFP in roughly 20 neurons at the L1 stage prior to remodeling [41]. Notably, in the ventral nerve cord, DVE-1 was solely expressed in DD GABAergic neurons (**Figure 5A,B**). Further, DVE-1 was specifically localized to GABAergic nuclei, where it assembled in discrete nuclear foci during the time frame of synaptic remodeling. Similar nuclear DVE-1 clusters have been observed in intestinal cell nuclei where DVE-1 is thought to regulate gene expression during the mitochondrial unfolded protein response (mtUPR) by associating with loose regions of chromatin and organizing chromatin loops [40]. DVE-1 localization to GABAergic nuclei raises the possibility that DVE-1 mediates its effects by regulating gene programs required for synapse elimination in GABAergic neurons. DVE-1::GFP expression in GABAergic neurons required the Pitx family homeodomain transcription factor UNC-30, the terminal selector of *C. elegans* GABAergic motor neuron identity (**Figure S5.1A**) [42–44]. Consistent with this observation, putative UNC-30 binding sites [42–44] are present within the DVE-1 promoter region. Interestingly, mutation of *unc-30* produced no appreciable change in DVE-1::GFP fluorescence in intestinal cells, indicating cell type-specific mechanisms for *dve-1* expression mediated at least in part through UNC-30 regulation (**Figure S5.1B**).

**Figure 5.**
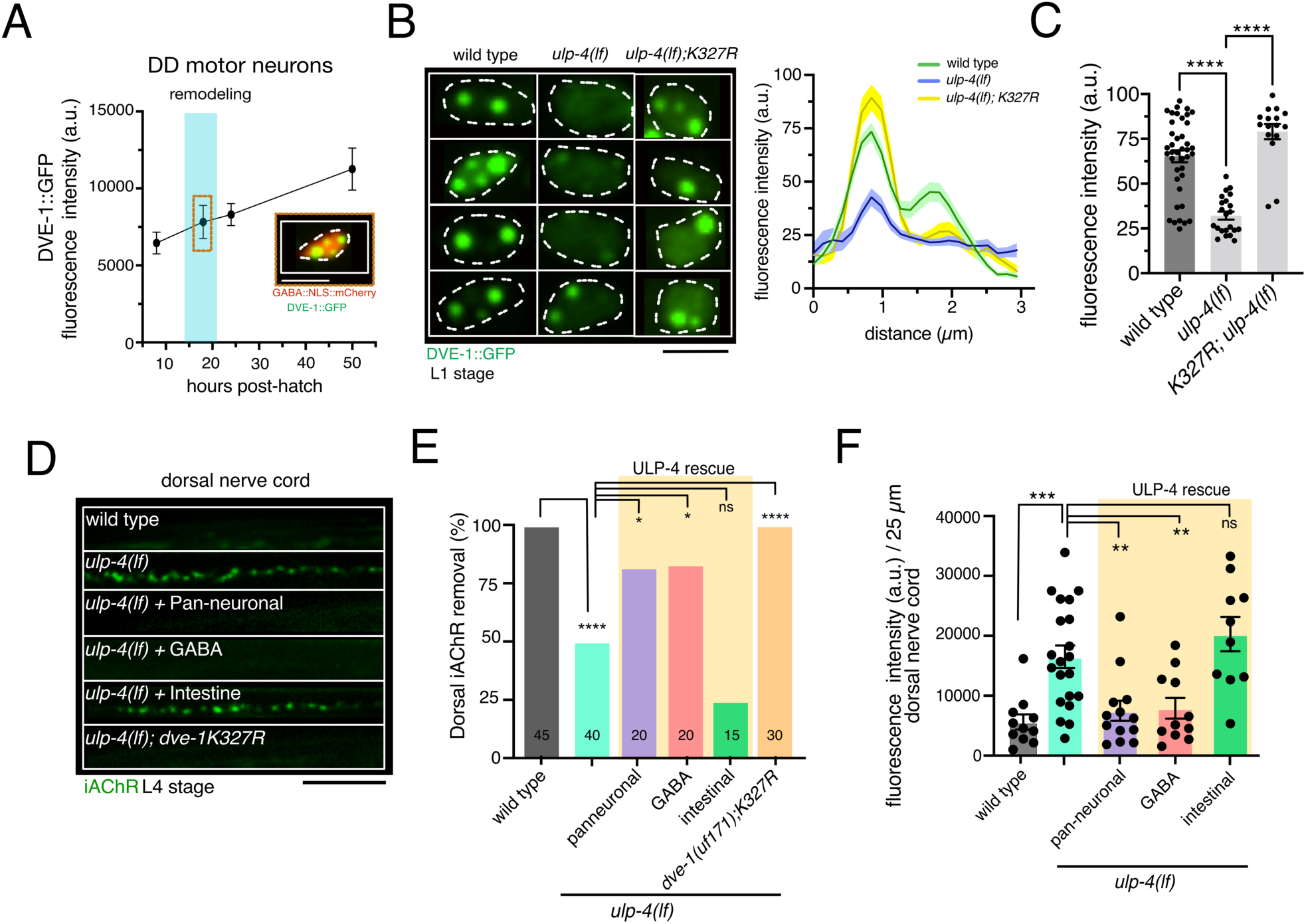
DeSUMOylating peptidase ULP-4 regulates nuclear localization of DVE-1. (A) Fluorescence intensity of nuclear DVE-1::GFP in DD motor neurons at 10, 18, 22 and 50 hours after hatch. Nuclear DVE-1::GFP is organized in discrete foci and increases through development. Each time point indicates mean ± SEM from at least three independent experiments. Inset, representative image of nuclear DVE-1::GFP in DD motor neuron 18 hours after hatch (nucleus labeled by NLS::mCherry). White dashed line outlines nucleus. Scale bar, 3 µm. (B) Confocal fluorescence images (left) of DVE-1::GFP in DD GABAergic motor neurons of L1 stage wild type, *ulp-4(lf)* mutant, and *K327R;ulp-4(lf)* double mutants. *ulp-4(lf)* disrupts DVE- 1::GFP nuclear localization. DVE-1(K327R) disrupts SUMOylation of DVE-1 and restores DVE-1::GFP nuclear localization in *ulp-4(lf)*. White dashed line outlines nucleus. Scale bar, 3 µm. Right, line scan of nuclear DVE-1::GFP fluorescence intensity in DD motor neurons of wild type (green) n=16, *ulp-4(lf)* (blue) n=11 and DVE-1(K327R)*;ulp-4(lf)* double mutants (yellow) n=8. Solid line represents mean, shading represents standard deviation of fluorescence. (C) Scatterplot (left) of the peak nuclear DVE-1::GFP fluorescence intensity in DD motor neurons. Each point represents a single DD nucleus. Imaged at least 2 DD cells per animal at L1 stage. Wild type: n=16, *ulp-4(lf)*: n=11, *K327R;ulp-4(lf)*: n=8. Bars indicate mean ± SEM. \*\*\*\**p*<0.0001, ANOVA with Dunnett’s multiple comparisons test. (D) Fluorescent confocal images of iAChR clusters in the dorsal processes of L4 stage DD neurons. iAChRs remained present in the dorsal nerve cord of L4 stage *ulp-4(lf)* mutants, indicating a failure in juvenile synapse elimination. Either pan-neuronal or specific expression of wild type *ulp-4* cDNA in GABAergic neurons was sufficient to restore synapse elimination. Scale bar, 5 µm. (E) Quantification of iAChR clustering at L4 stage for the genotypes indicated. Bars indicate the percentage of L4 stage animals where dorsal iAChRs have been completely removed. \**p*<0.05, \*\*\*\**p*<0.0001, ns – not significant, Fischer’s exact test with Bonferroni correction. 1/2 pan-neuronal lines, 2/2 GABA lines, and 0/2 intestinal rescue lines restored proper removal of dorsal iAChRs by L4. (F) Scatterplot of average iAChR fluorescence intensity/25 µm in the dorsal nerve cord at L4 stage for the genotypes indicated. Each point represents a single animal. Bars indicate mean ± SEM. \*\**p*<0.05, \*\*\**p*<0.001, ns – not significant, ANOVA with Dunnett’s multiple comparisons test. The number of animals was >10 per genotype.

Prior studies of DVE-1 in intestinal cells showed that deSUMOylation of DVE-1, mediated by the isopeptidase ULP-4, is required for its nuclear localization [45]. To explore whether DVE-1 localization to DD GABAergic nuclei is important for DVE-1 regulation of synapse elimination, we asked if the nuclear localization of DVE-1 in GABAergic neurons is also regulated by ULP-4. We observed that mutation of *ulp-4* caused a striking decrease of nuclear *dve-1::*GFP fluorescence in GABAergic neurons and severely diminished *dve-1::*GFP nuclear foci (**Figure 5B,C**). A mutated form of DVE-1::GFP, DVE-1K327R, where a key lysine residue required for SUMOylation is mutated to arginine [45], localized to GABAergic nuclei in the absence of *ulp-4* (**Figure 5B,C**). Together, these data demonstrate that nuclear localization of DVE-1 in GABAergic neurons is regulated by ULP-4 and SUMOylation. Importantly, we found that ULP-4 was also required for the elimination of dorsal iAChR clusters during remodeling, such that dorsal iAChR clusters remained present in roughly 50% of L4 stage *ulp-4* mutant animals (**Figure 5D-F**). Either pan-neuronal or GABA neuron-specific expression of the wild type *ulp-4* gene in *ulp-4* mutants was sufficient to restore the elimination of dorsal iAChRs, while intestinal *ulp-4* expression was not (**Figure 5D-F**). Further, mutation of the DVE-1 SUMOylation site (K327R) by itself did not impair synapse elimination but restored proper removal of iAChRs in *ulp-4* mutants (**Figure 5D-E**). We conclude that the localization of DVE-1 to GABAergic nuclei is essential for synapse elimination during remodeling, and this localization is regulated at least in part by SUMOylation. Notably, the nuclear localization of mammalian SATB family members is also dependent on SUMOylation, suggesting conserved regulatory mechanisms [46, 47].

### Analysis of DVE-1 transcriptional targets reveals several pathways with relevance for synapse elimination

Recent work revealed that homeodomain transcription factors are broadly utilized in the specification of *C. elegans* neuronal identity [41]. Given this finding and DVE-1 homology with mammalian SATB family transcription factors, we asked whether DVE-1 transcriptional regulation may be important for GABAergic neuronal identity. We found that the numbers of DD neurons and commissures were unchanged in *dve-1* mutants compared to wild type (**Figure S6.1A**). In addition, we found that the expression levels for *oig-1* (**Figure 4A**) and three additional GABAergic markers in DD neurons (*unc-47*/GABA vesicular transporter, *unc-25*/glutamic acid decarboxylase, and *flp-13*/FMRFamide neuropeptide) were not appreciably altered by *dve-1* mutation (**Figure S6.1B**). These results support that DVE-1 is not critical for GABAergic identity of the DD neurons but instead regulates other aspects of GABAergic neuron development, such as the transcription of effectors important for synapse elimination.

To reveal potential direct targets of DVE-1 in GABAergic neurons, we analyzed chromatin immunoprecipitation followed by sequencing (ChIP-Seq) data available from the modENCODE consortium [48]. We found 1044 genes with strong DVE-1 binding signal in their promoter regions, implicating these genes as potential direct targets of DVE-1 transcriptional regulation (**File S5**). We noted that 627 of these genes are significantly expressed in GABAergic neurons based on available single-cell RNA-seq data (**File S5**) [49, 50]. Pathway analysis of the GABAergic neuron-enriched targets using WormCat [51] and WormenrichR [52] revealed a significant enrichment of genes involved in the mitochondrial unfolded protein response (mtUPR) stress pathway (**Figure 6A, File S5)**, as expected from prior studies [27, 45, 53]. Notably, our analysis also revealed enrichment of genes involved in the ubiquitin-proteasome system as well as various other processes including ribosomal composition, and endocytic and phagocytotic function (**Figure 6A, B**, **Table 3 File S5**). These pathways represent intriguing potential targets for DVE-1-dependent regulation in the control of synapse elimination.

**Figure 6.**
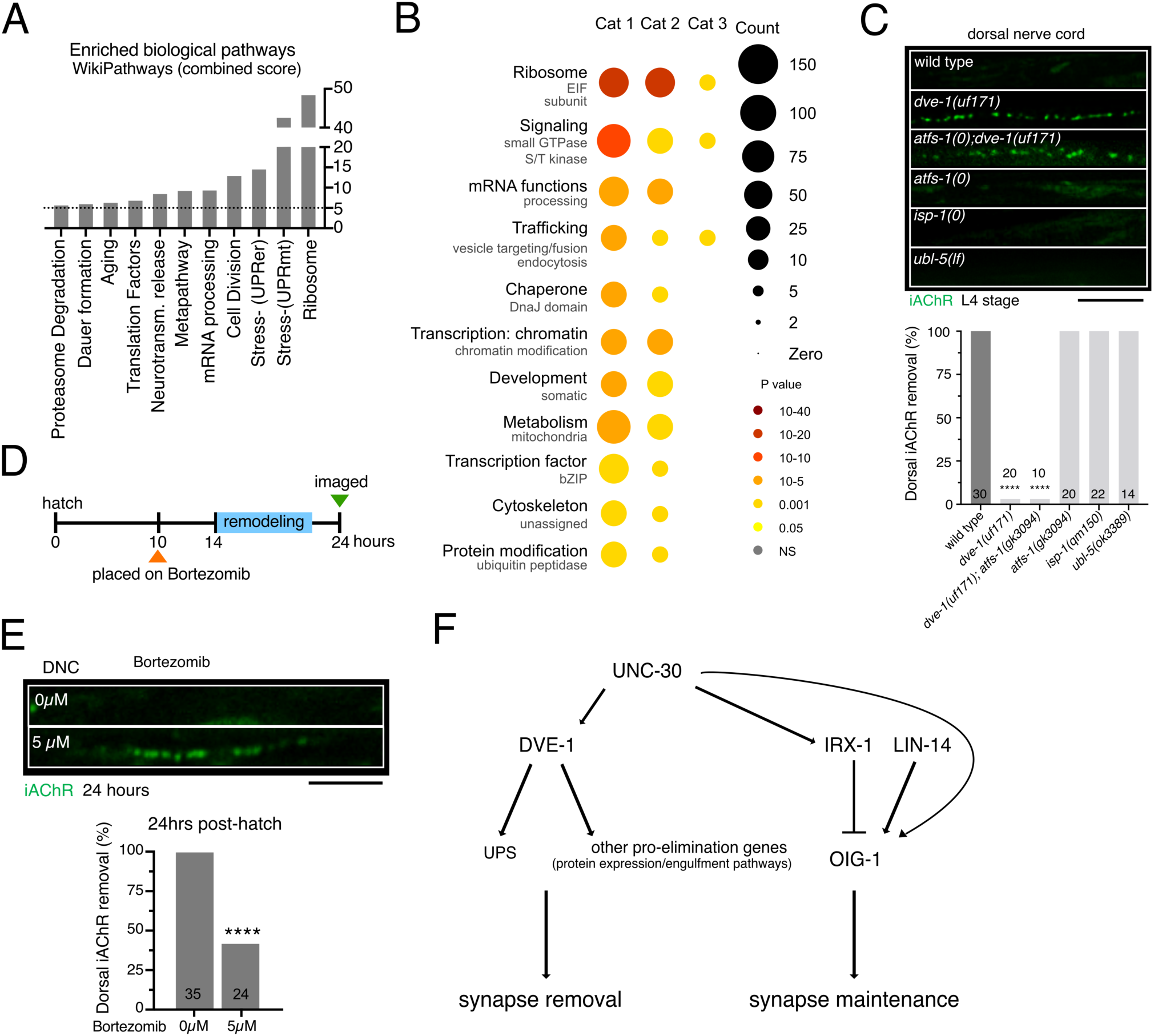
Enrichment analysis of DVE-1 ChIP-seq targets reveals potential pathways governing synapse elimination. (A) The WormEnrichR pathway enrichment analysis utilizing both the Kyoto Encyclopedia of Genes and Genomes (KEGG) database and the WikiPathway database. Bars represent enriched pathways with a combined score >10 (dashed line) and p-adj < 0.05. Y-axis is segmented between 40-60. (B) Wormcat analysis for significantly enriched categories of putative DVE-1 targets identified from ChIP-seq dataset. size of circles indicates the number of genes and color indicates the value of significant over-representation in each Wormcat category. (C) Induction or inhibition of mtUPR has no effect on synapse elimination. Top, fluorescent confocal images of iAChR clusters in the dorsal processes of DD neurons at L4 stage. Scale bar, 5 µm. Bottom, bars indicate the percentage of L4 stage animals that have eliminated dorsal iAChRs. \*\*\*\**p*< 0.0001, Fischer’s exact test with Bonferroni correction. (D) Schematic of Bortezomib inhibitor experimental design. Animals were placed on Bortezomib plates at 10 hours after hatch. animals were allowed to grow on this inhibitor until imaging at 24 hours after hatch. (E) Top, fluorescent confocal images of iAChR clusters in the dorsal processes of DD neurons at 24 hours post-hatch grown on (5µM) or off (0µM) Bortezomib inhibitor plates. Bottom, bars indicate the percentage of animals that have removed dorsal iAChRs by 24 hours post-hatch. \*\*\*\**p*< 0.0001, Fischer’s exact test. (F) Transcriptional control of DD remodeling. The GABAergic terminal selector transcription factor UNC-30/Pitx regulates the cellular expression of both *dve-1* and *oig-1* [23, 24], this paper. *oig-1* encodes an Ig-domain protein that stabilizes juvenile synapses prior to remodeling. Temporal control of *oig-1* expression is encoded through additional transcriptional regulation by LIN-14 and IRX-1 transcription factors [23, 24]. We show that DVE-1 promotes synapse removal/destabilization, perhaps through transcriptional regulation of the ubiquitin proteasome system (UPS). Mutation of *dve-1* impairs synapse removal, even when OIG-1 mediated stabilization is removed.

**Table 3:**
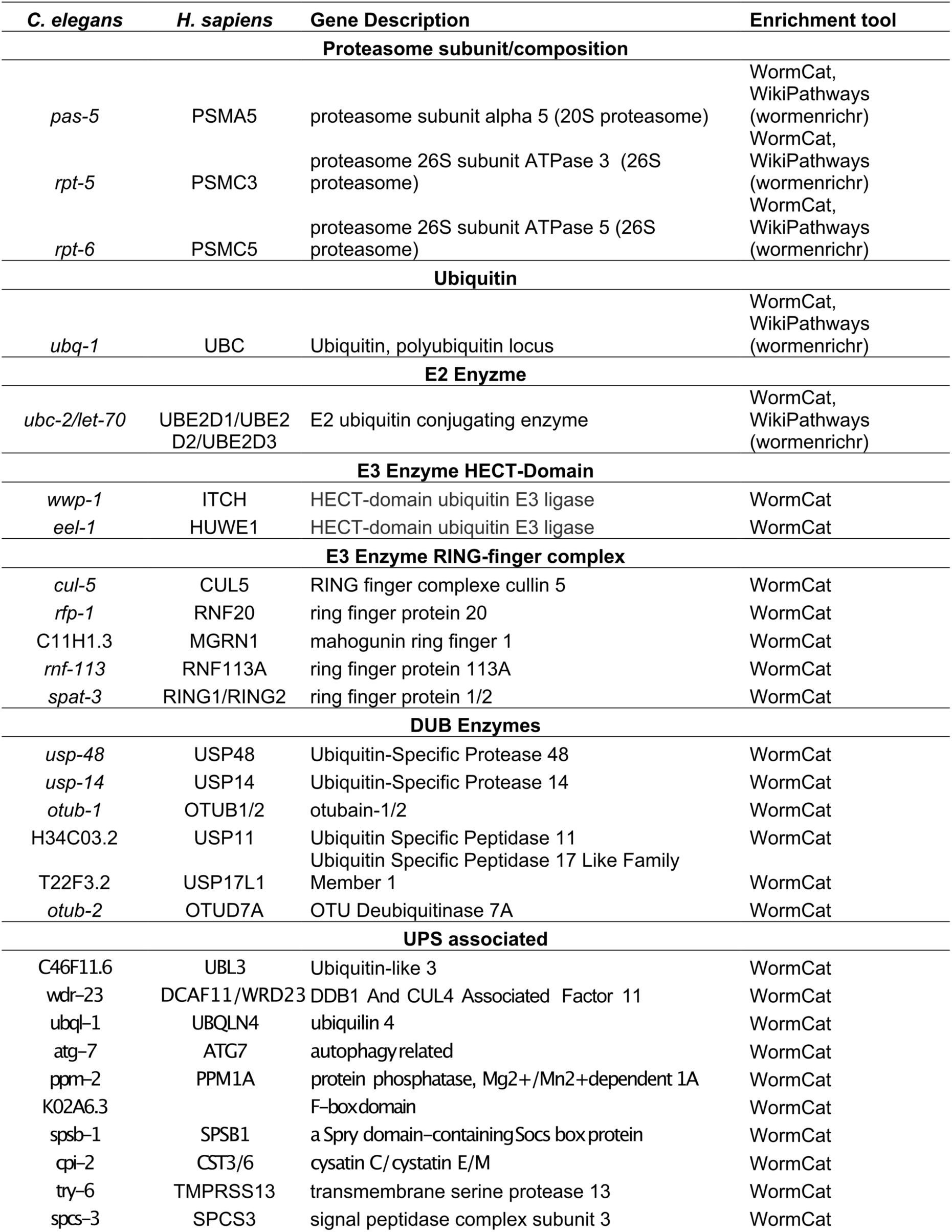
Potential ubiquitin proteasome system targets of DVE-1 revealed by ChIP-seq analysis

### Inhibition of the ubiquitin-proteasome system, but not activation or inhibition of mtUPR, delays DVE-1-dependent synapse elimination

To clarify which of the pathways identified from our analysis may be most critical, we next asked whether DVE-1 regulates synapse elimination by activating or inhibiting the mtUPR. We first quantified the length and density of mitochondria in DD neurons and found no differences between wild type and *dve-1* mutants (**Figure S6.2A**). We next measured mtUPR activation in *dve-1* mutants by quantifying the fluorescence of *hsp-6pr::*GFP, a commonly used mtUPR reporter [27]. Surprisingly, we noted increased levels of intestinal *hsp-6pr::*GFP expression in *dve- 1* mutants compared with control, suggesting elevated mtUPR activity (**Figure S6.2B, C**). The transcription factor ATFS-1 is required for the expression of *hsp-6pr::*GFP transcriptional response and activation of the mtUPR [54]. RNAi knockdown of *atfs-1* decreased *hsp-6pr::*GFP expression in *dve-1* mutants (**Figure S6.2C**), indicating that downregulation of *atfs-1* reduced the mtUPR. However, inhibition of the mtUPR by *atfs-1* knockdown failed to restore normal removal of dorsal iAChR clusters in *dve-1* mutants (**Figure S6.2D**). Likewise, a null mutation in *atfs-1* did not alter synapse elimination in otherwise wild type animals and failed to restore synapse elimination when combined with mutation of *dve-1* in *atfs-1;dve-1* double mutants (**Figure 6C**). These results show that increased activation of the mtUPR in *dve-1* mutants is not sufficient to account for a failure in synapse removal. Consistent with this interpretation, constitutive mtUPR activation by mutation of the mitochondrial complex III subunit gene *isp-1* also did not alter synapse elimination (**Figures 6C** **and S6.2D**) [55]. Additionally, mutation of *ubl-5*, a cofactor with DVE-1 in the initiation of the intestinal mtUPR [56], did not affect synapse elimination (**Figure 6C**). Our findings demonstrate that mtUPR activation or inhibition do not alter synapse removal. We conclude that DVE-1 coordinates synapse elimination through transcriptional regulation of alternate pathways, perhaps those identified from our enrichment analysis (**Figure 6A,B**).

Given recent evidence for the regulation of synapse structure through ubiquitin-dependent degradation processes and links with neurological disease [57], we next asked whether DVE-1 control of synapse elimination may occur through transcriptional regulation of ubiquitin-proteasome function. We investigated the requirement for the ubiquitin proteasome system (UPS) during synapse elimination using bortezomib, a small molecule inhibitor of the 26S proteasome. While treatment with high (≥10 µM) concentrations of bortezomib produced larval arrest, treatment with 5 µM bortezomib was sufficient to disrupt UPS function, as assessed by induction of the *skn-1* dependent proteasome reporter *rpt-3pr::*GFP [58] (**Figure S6.2E**), without causing developmental arrest. Moreover, treatment with 5 µM bortezomib significantly delayed synapse elimination during remodeling. Specifically, more than 50% of animals treated with bortezomib failed to remove juvenile dorsal postsynaptic sites in DD neurons by 24 hrs after hatch (**Figure 6D-F**), demonstrating a requirement for UPS function in synapse elimination. The identification of UPS pathway genes as targets for transcriptional regulation by DVE-1 therefore leads us to propose that DVE-1 control of synapse elimination during remodeling occurs at least in part through transcriptional regulation of the ubiquitin proteasome system in DD GABAergic neurons (**Figure 6G**).

## Discussion

Developmental remodeling of synaptic connectivity occurs throughout phylogeny, refining and reorganizing neuronal connections toward the establishment of the mature nervous system. While neuron-extrinsic events that shape remodeling, for example, microglial phagocytosis of synaptic material [5–8], have gained a lot of recent attention, neuron-intrinsic processes governing remodeling have remained less well-described. Likewise, the relationship between degenerative and growth processes during remodeling have not been clearly elucidated. The developmental remodeling of *C. elegans* GABAergic DD neurons presents a uniquely accessible system for addressing important questions about evolutionarily conserved neuron-intrinsic mechanisms of remodeling because the reorganization of their connectivity occurs without gross morphological changes or a requirement for synaptic removal by other cell types.

Here, we show that the homeodomain transcription factor *dve-1* with homology to the SATB family of transcription factors is specifically required for the elimination of juvenile synaptic inputs to DD neurons during remodeling. In *dve-1* mutants, juvenile postsynaptic sites and apposing cholinergic presynaptic terminals are preserved into adulthood. The failure to eliminate these sites results in elevated activity at these synapses and impaired motor function. Interestingly, *dve-1* is not required for the growth of new DD neuron synaptic inputs that are characteristic of the mature motor circuit, indicating that the formation of new connections is not dependent upon elimination of pre-existing juvenile synapses. Likewise, mutation of *dve-1* does not alter developmental reorganization of synaptic outputs from DD neurons onto muscles. In *dve-1* mutants, newly relocated GABAergic synaptic terminals occupy similar territories in DD neurons as lingering juvenile synaptic inputs. Thus, the formation of new GABAergic presynaptic terminals during maturation of the circuit is not contingent on elimination of nearby juvenile postsynaptic sites in DD neurons.

Our findings lead us to propose that cell-autonomous transcriptional regulation of GABAergic neurons by DVE-1 promotes the elimination of their juvenile synaptic inputs. We found that DVE- 1 is expressed in a limited number of neurons, including GABAergic motor neurons, and DVE-1 localization to GABAergic nuclei is required for synapse elimination to proceed normally. DVE-1 regulation of synapse elimination shares interesting parallels with a previously described pathway for elimination of postsynaptic structures at mouse glutamatergic synapses through transcriptional regulation by Myocyte Enhancer Factor 2 (Mef2) [59, 60]. However, in contrast to MEF2-regulated synapse elimination, we found that DVE-1-dependent elimination is not strongly activity-dependent. Also, we did not observe strong temporal regulation of DVE-1 expression in GABAergic neurons prior to or during remodeling, raising important mechanistic questions about the timing of synapse elimination. One possible route for temporal regulation might be through control of DVE-1 nuclear localization. We show that DVE-1 localization to GABAergic nuclei can be regulated through SUMOylation. However, we observed nuclear localization of DVE-1 in GABAergic neurons prior to the onset of remodeling, suggesting the presence of additional mechanisms for temporal control. This is consistent with prior work indicating that temporally controlled expression of OIG-1 regulates the timing of remodeling [23, 24]. We found that precocious synapse elimination in *oig-1* mutants is reversed when *dve-1* function is also disrupted, further indicating that DVE-1 transcriptional regulation is required for synapse elimination to occur. In contrast, precocious growth of new synapses in *oig-1* mutants was not altered by mutation of *dve-1*, suggesting that DVE-1 regulated degenerative mechanisms act in parallel with growth processes that are regulated independently (**Figure 6G**).

Our experiments show that mutation of *dve-1* affects the stability of both postsynaptic sites in DD GABAergic neurons and presynaptic terminals in cholinergic neurons. We show that the juvenile synaptic vesicle assemblies in axons of presynaptic cholinergic neurons are almost completely exchanged during the 10-hour period of wild type remodeling. This turnover of synaptic vesicles during remodeling contrasts with the relative stability of wild type synaptic vesicles over the 10 hours immediately following remodeling. Thus, turnover of synapse-associated vesicles in presynaptic cholinergic axons occurs at a higher rate during circuit rewiring, suggesting temporal regulation of an active process for presynaptic reorganization. Notably, removal of juvenile synaptic vesicle assemblies during remodeling is strikingly reduced in *dve-1* mutants, indicating that disruption of DVE-1 transcriptional activity is sufficient to stabilize presynaptic vesicle pools in cholinergic neurons. Since DVE-1 is solely expressed in postsynaptic GABAergic neurons, our results suggest that DVE-1-regulated postsynaptic pathways promote exchange or elimination of juvenile presynaptic elements through destabilization of the postsynaptic apparatus. Our photoconversion experiments also show that recruitment of new synaptic vesicles in cholinergic axons of *dve-1* mutants is not halted by the stabilization of juvenile synaptic vesicle assemblies. We noted an overall increase in dorsal synaptic vesicle material in *dve-1* mutants compared with either wild type or ventral synapses of *dve-1* mutants. Our Ca^2+^ imaging and behavioral experiments provide evidence that the increase in cholinergic synaptic vesicles of dorsally projecting motor neurons alters circuit function such that cholinergic activation of dorsal musculature is enhanced in *dve-1* mutants, resulting in deeper dorsal bends and a dorsal turning bias during movement.

Our pathway analysis of DVE-1 ChIP-seq data showed a high enrichment of genes involved in the mtUPR. In the mtUPR, DVE-1/SATB is thought to organize loose chromatin to induce expression of chaperones and proteases [27, 53, 56]. However, manipulations that either activated or inhibited the mtUPR did not affect remodeling, providing support for a model where DVE-1 regulation of remodeling is distinct from DVE-1 function in mtUPR. In addition to the mtUPR, our analysis of DVE-1 targets revealed enrichment of genes in other pathways that may represent alternative targets of DVE-1 for removal of synapses, in particular UPS pathway genes. Notably, inhibition of proteasome function caused a striking reduction in synapse elimination, supporting involvement of this pathway and suggesting that DVE-1 transcriptional regulation of the proteasome may be important to promote synapse elimination.

The closest homolog of DVE-1 is the *Drosophila* homeodomain transcription factor *defective proventriculus* (*Dve*). Interestingly, transcriptional profiling of *Drosophila* mushroom body gamma neurons during their remodeling showed Dve expression peaks at the onset of remodeling (https://www.weizmann.ac.il/mcb/Schuldiner/resources) [61]. DVE-1 also shares homology with the mammalian chromatin organizers special AT-rich sequence-binding (SATB) proteins 1 and 2. The SATB family of transcriptional regulators have been shown to play roles in many areas of mammalian brain development, such as the activation of immediate early genes important for maintaining dendritic spines in GABAergic interneurons [62] and cortex development and maturation [63], but roles in synapse elimination had been previously uncharacterized. Our findings offer a striking example of DVE-1/SATB transcriptional activation of pro-degenerative pathways acting in concert with temporally controlled expression of a maintenance factor to control a developmentally defined period of synapse elimination. Given the conservation of DVE- 1 and the mammalian SATB family of transcriptional regulators, our analysis may point toward new mechanisms by which SATB family transcription factors control brain development. Importantly, dysfunction of these transcription factors in humans, as in SATB2-associated syndrome, is characterized by significant neurodevelopmental delays, limitations in speech, and severe intellectual disability [64, 65] . More broadly, our findings highlight a cellular strategy for temporal control of circuit development through convergent regulation of antagonistic cellular processes. Interestingly, spatiotemporal regulation through competing parallel transcriptional programs is utilized in other developmental contexts across different species [66–68], suggesting this represents a broadly utilized mechanism for temporal control of key developmental events.

## Methods

### Strains and Genetics

All strains are derivatives of the N2 Bristol strain (wild type) and maintained under standard conditions at 20-25°C on nematode growth media plates (NGM) seeded with *E. coli* strain OP50. Some strains were provided by the *Caenorhabditis* Genetics Center, which is funded by NIH Office of Research Infrastructure Programs (P40 OD010440), and by the National BioResource Project which is funded by the Japanese government. Transgenic strains were generated by microinjection of plasmids or PCR products into the gonad of young hermaphrodites. Integrated lines were produced by X-ray irradiation or UV-integration and outcrossed to wild type/N2 Bristol. A complete list of all strains used in this work is included in supplemental file 1.

### Molecular Biology

Plasmids were constructed using two-slot Gateway Cloning system (Invitrogen) and confirmed by restriction digest and/or sequencing as appropriate. All plasmids and primers used in this work are described in supplemental file 2 and 3 respectively.

To generate the DVE-1 genomic rescue the *dve-1* promoter (5 kb upstream from translational start site), genomic fragment (from translational start to stop, 6107bp), and *dve-1* 3’ UTR (626bp downstream from translational stop) were amplified from genomic N2 DNA. pKA110 (*unc- 129pr*::Dendra2::RAB-3) was created by ligating a gBlock (IDT) containing the Dendra2 coding sequence into NheI-HF/PstI-HF digested pDest-114 to generate pDest-339 (Dendra2::RAB-3). pDest-339 was then recombined with pENTR-*unc-129* to generate pKA110 and injected at 50 ng/µl. To generate pKA35 (*unc-129pr*::Chrimson::SL2::BFP) Chrimson was amplified from pDest-104 and ligated into NheI-HF/BstBI digested pDest-239 to generate pDest-240 (Chrimson::SL2::BFP). pDest-240 was then recombined with pENTR-*unc-129* to generate pKA35 and injected at 50 ng/µl.

To generate ULP-4 rescue constructs ULP-4 cDNA was amplified from RNA and ligated into NheI-HF/KpnI-HF digested pDest-139 to generate pDest-291(ULP-4cDNA). pDest-291 was recombined with pENTR-*unc-47* to generate pKA76 (*unc-47pr*::ULP-4cDNA), pENTR-F25B3.3 to generate pKA78 (F25B3.3pr::ULP-4cDNA), and pENTR-*gly-19* to generate pKA80 (*gly- 19pr::*ULP-4cDNA). All *ulp-4* rescue constructs were injected at 30 ng/µl.

pKA74 (*unc-47pr*::NLS::mCherry) was created by amplifying an artificial intron and NLS from plasmid #68120 (Addgene) and was ligated into AgeI-HF/XbaI digested pDest-145 to generate pDest-205(NLS::mCherry). pDest-205 was recombined with pENTR-*unc-47* to generate pKA74 and injected at 50 ng/µl.

### Staging time course DD remodeling

Briefly, freshly hatched larvae were transferred to seeded OP50 plates and maintained at 25°C (timepoint 0). Imaging and analysis of iAChR or synaptic vesicle remodeling was assessed as previously described [23].

### Confocal imaging and analysis

Unless noted otherwise, all strains were immobilized with sodium azide (0.3M) on a 2% or 5% agarose pad. All confocal Images were obtained either using an Olympus BX51WI spinning disk confocal equipped with a 63x objective or on the Yokogawa CSU-X1-A1N spinning disk confocal system (Perkin Elmer) equipped with EM-CCD camera (Hamamatsu, C9100-50) and 63x oil immersion objective. Analysis of synapse number and fluorescence intensity was performed using FIJI ImageJ software (open source) using defined threshold values acquired from control experiments for each fluorescent marker. Statistical analysis for all synaptic and spine analysis between two groups utilized a student’s t-test; for analysis where, multiple groups were compared an ANOVA with Dunnett’s multiple comparisons test was used.

### Synaptic analysis

Background fluorescence was subtracted by calculating the average intensity of each image in a region outside the ROI. All ROIs were 25 µm or 30 µm in length. Quantification of the number of puncta within an ROI had a set threshold of 25-255 and the analyze particles function of ImageJ was used to quantify any particle greater than 4 pixels^2^. Fluorescence intensity was quantified from the raw integrated fluorescence within the ROI. For quantification of the DD neuron synapses the ROI was defined as either the ventral region anterior to the DD1 soma or the opposing dorsal region. For quantification of the DA/DB neuron synapses the ROI was defined as either the ventral region between DB1 and DB3 or the opposing dorsal region between VB1 and VB3.

### Spine/dendrite analysis

Spine number was quantified as described previously [32, 33]. Briefly, spines were counted within a 30 µm ROI anterior to the soma of DD1. Dendrite length was defined as the anterior extension from DD1 soma to the end of the ventral process.

### EMS Screen

The EMS mutagenesis protocol was adapted from [69]. *flp-13pr::*ACR-12::GFP strain IZ1905 animals were treated with 25 µM ethyl methanesulfonate (EMS, Sigma). After washing, P0 mutagenized animals were recovered. F1 animals were transferred to fresh plates and 8 F2s were isolated per F1 (F2 clonal screen). The F3 progeny of ∼400 F2s per round were screened. After 27 rounds of EMS, a total of 3261 haploid genomes were screened. Each isolated candidate mutant was rescreened three times to confirm the phenotype.

### Variant discovery mapping and whole genome sequencing

Mutant strains were backcrossed a single time into the IZ2302 starting strain injected with *unc- 122pr::*GFP co-injection marker(enabling the distinction of cross- from self-progeny). F2 animals were isolated onto separate plates and their F3 brood were screened on a confocal microscope for a remodeling phenotype. 21 independent homozygous recombinant F2 lines were thus isolated and pooled together. Worm genomic DNA was prepared for sequencing following Gentra Puregene Tissue Kit DNA purification protocol (Qiagen). Library construction and whole genome sequencing were performed by Novogene. Briefly, NEBNext DNA Library Prep Kit was used for library construction. Pair-end sequencing was performed on Illumina sequencing platform with a read length of 150 bp at each end. Reads were mapped to *C. elegans* reference genome version WS220 and analyzed using the CloudMap pipeline [70] where mismatches were compared to the parental strain as well as to the other sequenced mutants isolated from this screen [70, 71].

### Injection of fluorescent antibodies for *in vivo* labeling of iAChRs

For staining of iAChRs at the cell surface, mouse monoclonal α-HA antibodies (16B12 Biolegend) coupled to Alexa594 were diluted in injection buffer (20 mM K_3_PO_4_, 3 mM K citrate, 2% PEG 6000, pH 7.5). Antibody was injected into the pseudocoelom of L2/L3 stage wild type or *dve-1(uf171)* animals as described previously [23, 34]. Animals were allowed to recover for six hours on seeded NGM plates. Only animals in which fluorescence was observed in coelomocytes were included in the analysis. A student’s t-test was used for statistical analysis.

### Photoconversion of Dendra2::RAB-3

Wild type and *dve-1(uf171)* mutant L1 animals (12-14 hours post-hatch) expressing Dendra2::RAB3 were paralyzed using 1 mM Levamisole and imaged. Dendra2::RAB-3 puncta in the DA/DB dorsal axonal process were photoconverted using a 405 nm laser at 800 ms for 30 s. Images were acquired immediately following photoconversion and again 10 hours later. Between photoconversion and next timepoint animals were rescued and allowed to recover. Both red and green fluorescent signals were quantified. A student’s t-test was used for statistical analysis.

### Aldicarb paralysis assays

The aldicarb assays were performed as previously described [39]. Strains were scored in parallel, with the researcher blinded to the genotype. Young adult animals (24 hours after L4) at room temperature (22–24°C) were selected (>10 per trial for at least 3 trials) and transferred to a nematode growth medium plate containing 1 mM aldicarb (ChemService). Movement was assessed every 15 minutes for 2 hours. Animals that displayed no movement when prodded (paralyzed) were noted. The percentage of paralyzed animals were calculated at each timepoint.

### Calcium imaging

Transgenic animals expressing *ttr-39pr*::GCaMP6s::SL2::mCherry (GABA neurons) along with *unc-129pr*::Chrimson::SL2::BFP (DA and DB cholinergic neurons) were grown on NGM plates with OP50 containing 2.75 mM All-Trans Retinal (ATR). L4 animals (40 hours post-hatch) were staged 24 hours prior to experiments on fresh ATR OP50 NGM plates. Imaging was performed with 1-day age adults immobilized in hydrogel [32, 72]. Animals were transferred to 7.5 μL of the hydrogel mix placed on a silanized glass slide and covered with cover slip. Hydrogel was cured using a handheld UV Transilluminator (312 nm, 3 minutes). Chrimson photoactivation (∼14 mW/cm^2^) was achieved using a TTL-controlled 625 nm light guide coupled LED (Mightex Systems). A 556 nm BrightLine single-edge short-pass dichroic beam splitter was positioned in the light path (Semrock) [33]. Data were acquired at 10 Hz for 15 s using Volocity software at a binning of 4x4. Analysis was performed using ImageJ. DD and VD GABA motor neuron cell bodies were identified by mCherry fluorescence and anatomically identified by position along the ventral nerve cord. Each field typically contained 1–5 GABA motor neurons. Only recordings of neurons located anterior to the vulva were included in the analysis. Photobleaching correction was performed on background subtracted images by fitting an exponential function to the data (CorrectBleach plugin, ImageJ). Pre-stimulus baseline fluorescence (F_0_) was calculated as the average of the corrected background-subtracted data points in the first 4 s of the recording and the corrected fluorescence data was normalized to prestimulus baseline as ΔF/F_0_, where ΔF=F– F_0_. Peak ΔF/F_0_ was determined by fitting a Gaussian function to the ΔF/F_0_ time sequence using Multi peak 2.0 (Igor Pro, WaveMetrics). All data collected were analyzed, including failures (no response to stimulation). Peak ΔF/F_0_ values were calculated from recordings of >10 animals per genotype. Mean peaks ± SEM were calculated from all peak ΔF/F_0_ data values. For all genotypes, control animals grown in the absence of ATR were imaged.

### Single worm tracking

Single worm tracking was carried out for a duration of 5 minutes, on NGM plates seeded with 50 µL of OP50 bacteria, using Worm Tracker 2 [73]. Animals were allowed to acclimate for 30 s prior to tracking. Movement features were extracted from 5 minutes of continuous locomotion tracking. Worm tracker software version 2.0.3.1, created by Eviatar Yemini and Tadas Jucikas (Schafer lab, MRC, Cambridge, UK), was used to analyze movement [74]. Locomotion paths were extracted from 5 minutes of continuous locomotion tracking. Scoring of path trajectories was performed blinded to genotype.

### Optogenetic analysis

Behavioral assays were performed with young adults at room temperature (22°C–24°C). Animals were grown on plates seeded with OP50 containing 2.7 mM All-Trans Retinal (ATR). Animals were placed on fresh plates seeded with a thin lawn of OP50 containing ATR and allowed to acclimate for 1 minute. Dorsal-ventral position was noted prior to recording. Animals were allowed to move freely on plates and recorded with a Mitex X camera for 1 minute before stimulus, following this a Mightex LED module was used to stimulate Chrimson (625nm 14mW/cm^2^) continuously for 2 minutes. Locomotion (trajectory and body bending) was recorded with WormLab (MBF Bioscience) software. A mid-point bending angle histogram was generated for each animal such that over the span of 2 minutes body angles were measured and binned by the degree of angle. Depending on starting position negative and positive degree angles were assigned dorsal or ventral. Any bending angle greater than 0 but less than 50° was determined a regular bend. We noted wild type animals without stimulus rarely make angles greater then 50° and qualified any bending angle over 50° as a deep bend. An ANOVA with Dunnett’s multiple comparisons test was used for comparisons between pre-stimulus and stimulus in wild type and *dve-1(uf171)*. A Student’s T-test was used when comparing number of dorsal bends greater then 50° in wildtype vs *dve-1* mutant animals.

### RNAi by feeding

L4 larvae expressing *hsp-6pr::*GFP were cultured with *E. coli* expressing either control double-stranded RNA (empty vector) or targeting *atfs-1* and progeny were allowed to develop to L4 stage at 20°C. Intestinal GFP fluorescence of L4 stage progeny was measured using the Zeiss Imager M1, 10x objective.

### CRISPR/Cas9 K327R

Strain IZ4473 *dve-1(uf206)* was generated in *syb1984* (DVE-1 CRISPR-Cas9-mediated GFP knock, Tian lab) animals. A K-R mutation was created by changing AAA to CGT in exon 6, 4783 bp downstream of start. The IDT CRISPR HDR design tool (https://www.idtdna.com/pages/tools/alt-r-crispr-hdr-design-tool) was used to generate repair templates and guide sequences. Animals were injected with CRISPR/Cas9 mix [crRNA (oligo 2nmol, IDT), Donor (oligo 4nmol, IDT), purified Alt-R S.p. Cas9 nuclease V3 100µg (IDT CAT 1081058), Alt-R CRISPR/Cas9 tracrRNA (5nmol, IDT CAT 1072532), and pRF-4 (*rol-6* plasmid)]. Rolling worms were singled and validated by PCR sequencing. CRISPR/Cas9 design is provided in supplemental file 4.

### DVE-1 nuclear localization

DVE-1::GFP was measured in L1 stage DD nuclei using the strain *syb1984* (DVE-1 CRISPR- Cas9-mediated GFP knock in); *ufEx1814*(*unc-47pr::NLS::mCherry*). ROIs were determined by expression of the nuclear localized mCherry signal. Within the ROI a segmented line was drawn through the nucleus and an intensity profile was created for each nucleus. Fluorescence intensity values for DVE-1::GFP were quantified by averaging the largest 5 intensity values at the peak (roughly 0.5 µm). At least 2 DD nuclei per animal were analyzed. ANOVA with Dunnett’s multiple comparison test was used for statistical analysis. For analysis of *unc-30* mutants, an ROI was selected from the base of the pharynx out posteriorly by 30µm. For consistent analysis red GABA neurons within the head, unaffected by *unc-30* mutation and the pharynx served as landmarks for both wild type and *unc-30* mutant animals. Students t-test was used for statistical analysis.

### ChIP-seq data acquisition from ModEncode

modENCODE (www.modencode.org) produced ChIP-seq data by immunoprecipitating GFP from a line expressing DVE-1::GFP that was stably integrated into the genome. The DVE-1 ChIP-seq data set included two biological replicates at the late embryo stage as well as control animals. Significant peaks were called using PeakSeq and only peaks that were identified in both biological replicates were considered for analysis. DVE-1::GFP ChIP-seq data and experimental details can be found at http://intermine.modencode.org/release-33/report.do?id=77000654 (DCCid: modENCODE_4804) [75, 76]. Peaks were considered mapped to genes if there was at least an 80% overlap between the peak maximum read density and the region within 1kb base pairs upstream of the genes annotated transcriptional start site using the UCSC table browser intersect function [77].

### Pathway analysis

Pathway analysis was performed using both WormCat [51]: http://www.wormcat.com, and the enrichment analysis tool WormenrichR [52, 78]: https://amp.pharm.mssm.edu/WormEnrichr/. For WormCat analysis pathways were considered enriched if they had a p-value < 0.01 and Bonferroni FDR < 0.01. The WormEnrichR pathway enrichment analysis utilized the WikiPathway database [79]. For WormEnrichR, pathways were considered enriched if they had a p-adj < 0.05 and a combined score >10.

### Bortezomib time course

Worms were hatched synchronously on NGM plates at 25°C. Animals were transferred to 5µM Bortezomib (MilliporeSignma) plates at 10 hours post-hatch and allowed to develop until imaging at 24 hours post-hatch.

## Acknowledgements

We would like to thank Alexandra Byrne, Dori Schafer and the members of the Francis lab for critical reading of the manuscript. We also thank Ye Tian and Cole Haynes for generously sharing reagents and Michael Gorczyca and Will Joyce for technical assistance. We would like to thank Mark Alkema and Jeremy Florman for assistance with single worm tracking and analysis scripts. Some nematode strains were provided by the *Caenorhabditis* Genetics Center which is funded by the NIH National Center for Research Resources and the National Bioresource Project Nematode of Japan.

