## Supplemental Figures for "The homeodomain transcriptional regulator DVE-1 directs a program for synapse elimination during circuit remodeling"

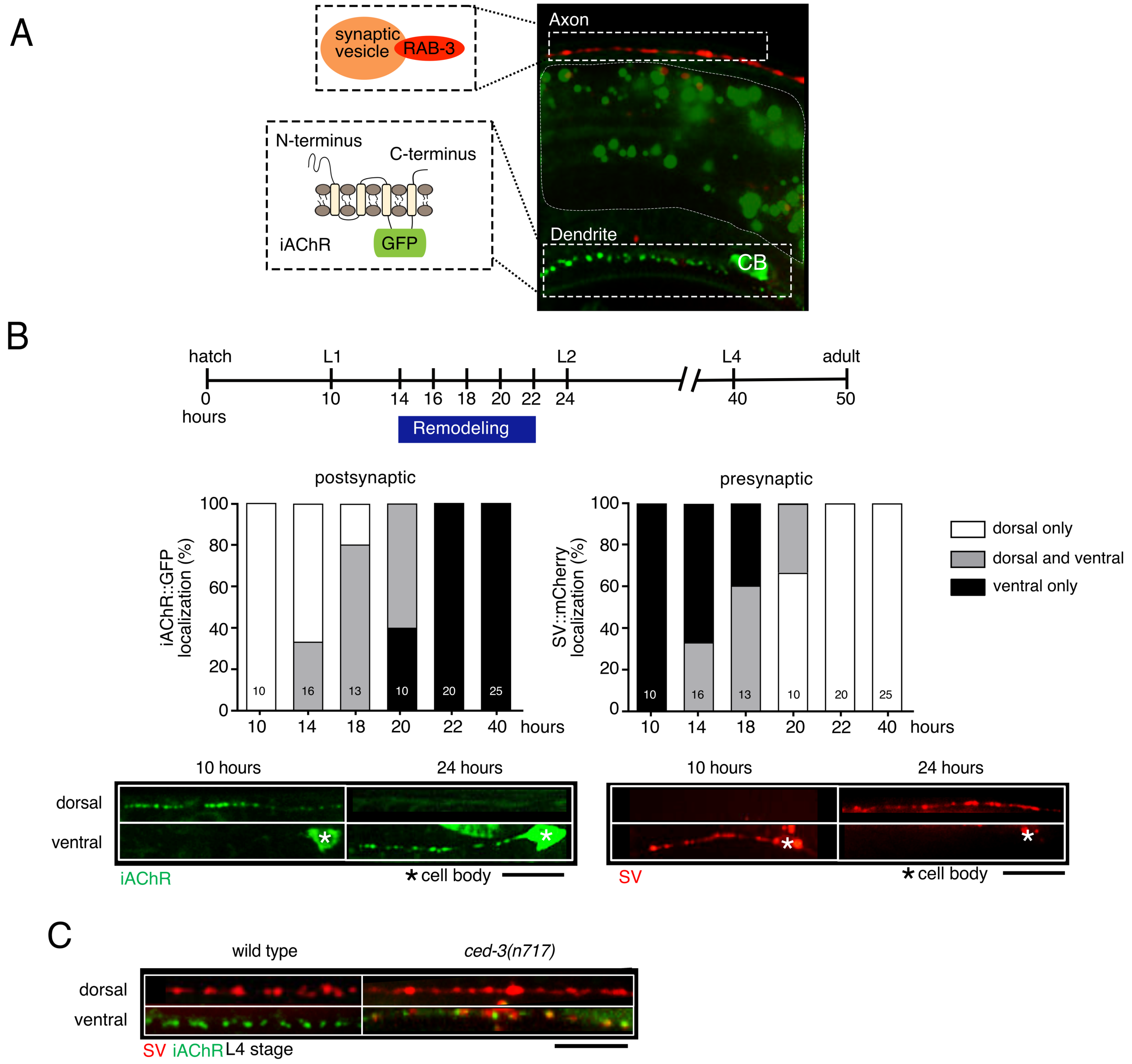


**Supplemental Figure 1.1. Remodeling of cholinergic postsynaptic sites and GABAergic presynaptic terminals occur simultaneously but are regulated through distinct mechanisms**

(A) Confocal image L4 stage wild type animal co-expressing a synaptic vesicle marker, defined in inset as mCherry::RAB-3, and iAChR marker, defined in inset as ACR-12::GFP in DD neurons. Insets are schematic representations of each marker. Masking of intestinal autofluorescence is outlined by small white dotted line.

(B) Top, timeline of wild type development. Approximate timing of transitions between larval stages and to adulthood are indicated. Blue bar indicates duration of DD synaptic remodeling in wild type animals. Middle, quantification of iAChR (postsynaptic) remodeling (left) and SV (presynaptic) remodeling in DD neurons (right) at the indicated time points after hatch. Animals are binned as dorsal only (white), ventral only (black), or dorsal and ventral (grey) according to the distribution of iAChR (left) or SV (right) clusters. Bottom, representative images of dorsal and ventral iAChR clusters (left) and SV puncta (right) at the times indicated in DD neurons of wild type animals. Remodeling of iAChR clusters and SV puncta occur simultaneously. Scale bar, 5 μm.

(C) Merged confocal images showing SV puncta (red) and iAChR clusters (green) in dorsal and ventral DD neuron processes of wild type (left) and *ced-3(n717)* mutants (right). Scale bar, 5 μm.

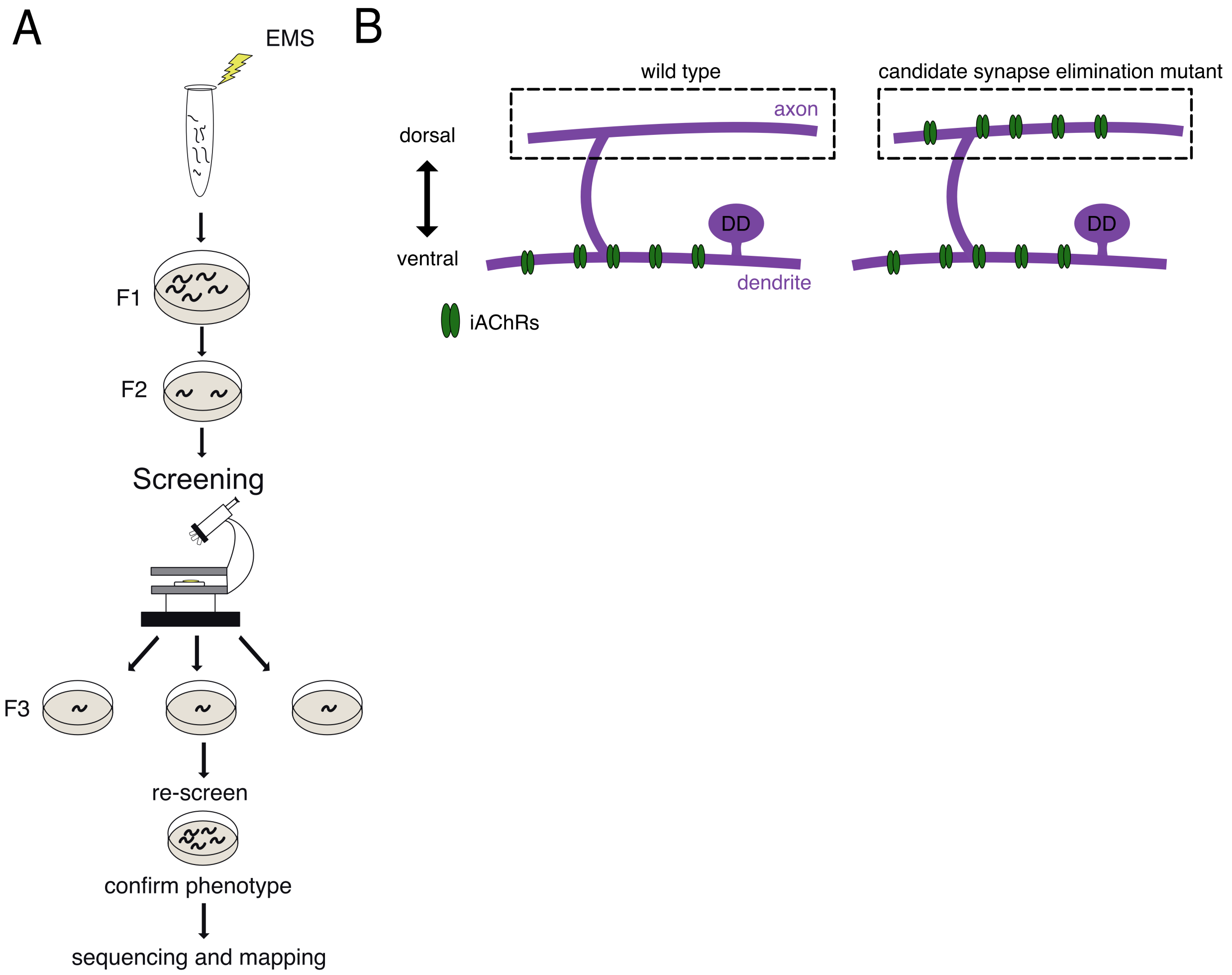


**Supplemental Figure 1.2. A Forward genetic screen to identify conserved mechanisms controlling synapse elimination**

(A) Schematic of experimental workflow for ethyl methanesulfonate (EMS) screen to obtain mutants with defects in the elimination of juvenile dorsal iAChR clusters.

(B) Schematics of iAChR localization within DD neurons of L4 stage wild type (left) or potential synapse elimination mutant (right).

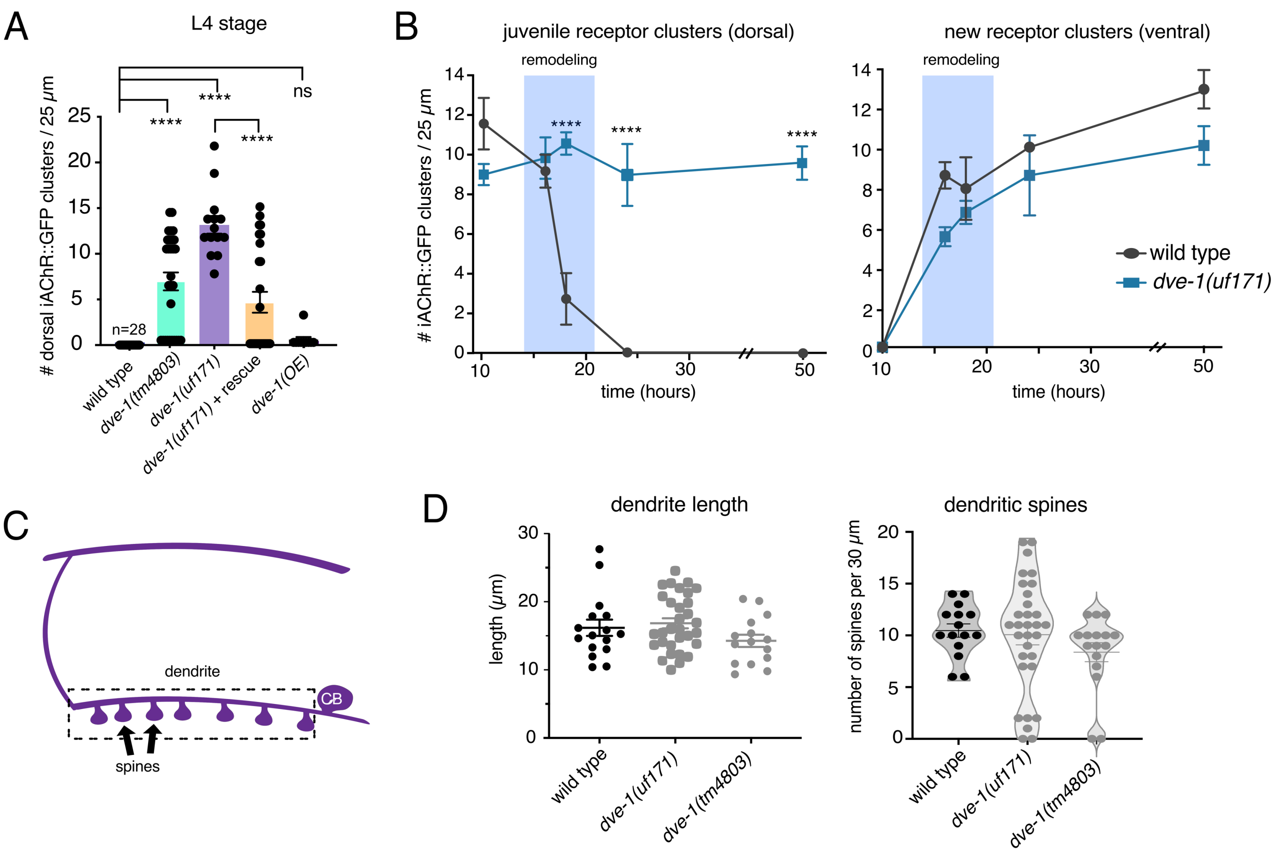


**Supplemental Figure 1.3. *dve-1* mutation disrupts synapse elimination but does not affect new synapse formation**

(A) Quantification of average number of iAChR clusters per 25 µm in the dorsal nerve cord of L4 stage DD neurons for the genotypes. Bars indicate mean ± SEM. ****$p$<0.0001, ns - not significant, ANOVA with Dunnett's multiple comparisons test. Each point represents a single animal. The number of animals >10 per genotype.

(B) Average number of juvenile iAChR clusters in the dorsal nerve cord (left) and new iAChR clusters in the ventral nerve cord (right) at indicated times after hatch. iAChR clusters are removed from the dorsal nerve cord of wild type animals (black) during remodeling (blue shading) but persist in in the dorsal nerve cord of *dve-1* mutants (blue). iAChR clusters in the ventral nerve cords of wild type and *dve-1* mutants increase similarly over time. Data points indicate mean ± SEM. ****$p<0.0001$, student's t-test. The number of animals >10 per genotype.

(C) Schematic of DD neuron, segmented box represents area quantified in D. Arrows indicate dendritic spines, CB, cell body.

(D) Scatterplot of average length of DD neuron dendrite (left) and number of dendritic spines (right) in wild type (purple), *dve-1(uf171)* (dark blue), *dve-1(tm4803)* (teal) overlayed with a violin plot to show distribution. Each point represents a single animal. The number of animals >10 per genotype.

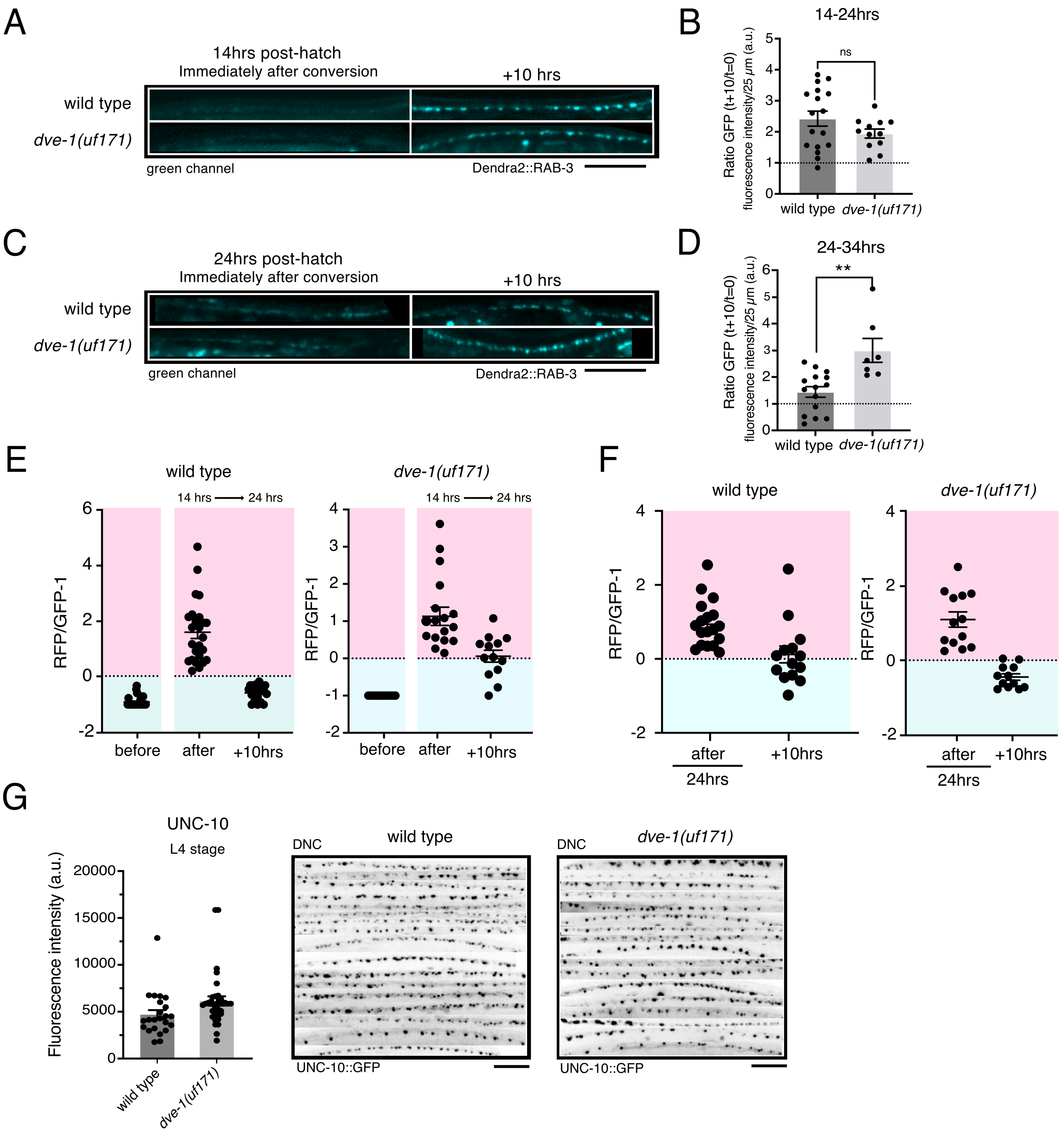


**Supplemental Figure 2.1. Presynaptic cholinergic synaptic vesicles turnover during remodeling**

(A) Confocal images of the dorsal nerve cord of wild type (top) and *dve-1(uf171)* mutants (bottom) showing green Dendra2::RAB-3 clusters in cholinergic DA/B neurons either immediately after photoconversion from green to red at 14 hours after hatch (left) or 10 hours later (right). Similar amounts of green Dendra2::RAB-3 clusters are added during remodeling of wild type and *dve-1* mutants. Scale bar, 5 µm.

(B) Scatterplot showing green Dendra2::RAB-3 fluorescence intensity 10 hours after photoconversion normalized to fluorescence intensity immediately after photoconversion prior to remodeling (10 hours after hatch) for wild type (left) and *dve-1(uf171)* mutants (right). Each point indicates a single animal. The number of animals >10 per genotype. Bars indicate mean ± SEM. ns - not significant, student's t-test.

(C) Confocal images of the dorsal nerve cord of wild type (top) and *dve-1(uf171)* mutants (bottom) showing green Dendra2::RAB-3 clusters in cholinergic DA/B neurons either immediately after photoconversion from green to red at 24 hours after hatch (left) or 10 hours later (right). Similar amounts of green Dendra2::RAB-3 clusters are added during remodeling of wild type and *dve-1* mutants. Scale bar, 5 µm.

(D) Scatterplot showing green Dendra2::RAB-3 fluorescence intensity 10 hours after photoconversion normalized to fluorescence intensity immediately after photoconversion at 24 hours after hatch (after remodeling) for wild type (left) and *dve-1(uf171)* mutants (right). Each point indicates a single animal. The number of animals >6 per genotype. Bars indicate mean ± SEM. ** $p<0.01$, student's t-test.

(E) Scatterplots of Dendra2-RAB-3 RFP/GFP fluorescence intensity ratio measurements before photoconversion at 10 hours post hatch (before remodeling), immediately after, and 10 hours later for wild type and *dve-1* mutants. Expressed as RFP/GFP fluorescence ratio -1 for display purposes. Values less than zero indicate enhanced green fluorescence while values greater than 1 indicate enhanced red fluorescence. Each dot represents a single animal. The number of animals > 10 per genotype/timepoint.

(F) Scatterplots of Dendra2-RAB-3 RFP/GFP fluorescence intensity ratio measurements before photoconversion at 24 hours after hatch (after remodeling), immediately after, and

10 hours later for wild type and *dve-1* mutants. Expressed as RFP/GFP fluorescence ratio-1 for display purposes. Values less than zero indicate enhanced green fluorescence while values greater than 1 indicate enhanced red fluorescence. Each dot represents a single animal. The number of animals > 10 per genotype/timepoint.

(G) Left, scatterplot showing average UNC-10::GFP fluorescence intensity in L4 stage cholinergic neurons (*acr-5pr*::UNC-10::GFP) of the dorsal nerve cord (DNC) for wild type and *dve-1* mutants. Each point indicates a single animal, the number of animals >10 per genotype. Bars indicate mean ± SEM. Right, stacked fluorescent images of the dorsal nerve cord showing UNC-10::GFP clusters in cholinergic neurons of L4 stage wild type and *dve-1(uf171)* mutants. Images on each line are from different animals. Scale bar, 5 μm.

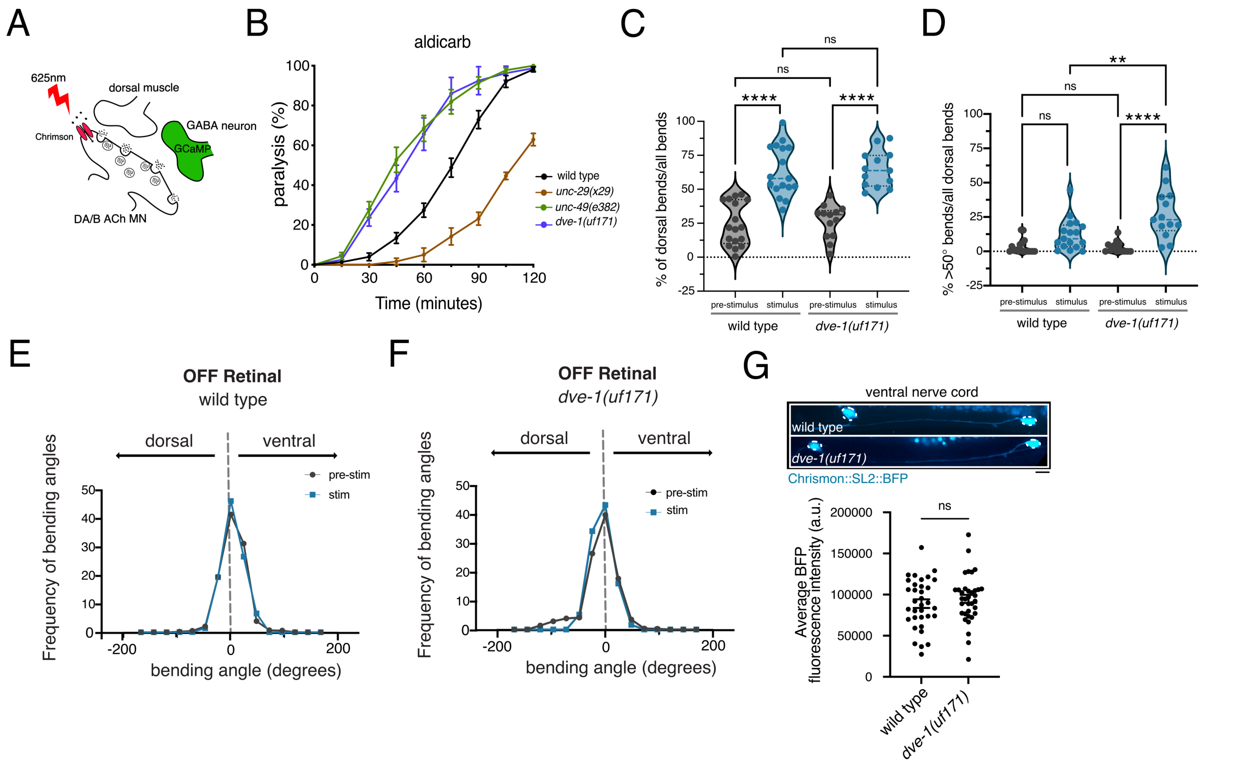


**Supplemental Figure 3.1. Motor circuit function is disrupted in *dve-1* mutants**

(A) Schematic of combined cell-specific expression of Chrimson (red receptor) for cholinergic depolarization, and GCaMP6s monitoring [$Ca^{2+}$] changes in the post-synaptic GABAergic motor neurons (green).

(B) Time course of paralysis in the presence of aldicarb (1 mM) for wild type (black) (n=14), *unc-29(x29)* mutants (brown) (n=6), *unc-49(e382)* mutants (dark green) (n=16), *dve-1(uf171)* mutants (blue) (n=12), are shown. At least 10 animals per trial. Data represent mean ± SEM.

(C) Scatterplot with violin overlay of the percentage of dorsal bends for wild type and *dve-1(uf171)* mutants before and after photostimulation. Each point represents a single animal. The number of animals > 10 per genotype. ****$p<0.0001$, ns - not significant, ANOVA with Dunnett’s multiple comparisons test.

(D) Scatterplot with violin overlay of the percentage of dorsal turns greater than 50° wild type (black) and *dve-1(uf171)* (blue) before and after photostimulation. Each point represents a single animal. The number of animals > 10 per genotype. **$p<0.01$, ****$p<0.0001$, ns - not significant, ANOVA with Dunnett's multiple comparisons test.

(E) Frequency distribution of body bending angles prior to (black) and during photostimulation (blue) for control animals in the absence of all-trans-retinal. Negative bending angle values indicate dorsal, while positive bending angle values indicate ventral. n=3.

(F) Frequency distribution of body bending angles prior to (black) and during photostimulation (blue) for *dve-1* mutants in the absence of all-trans-retinal. Negative bending angle values indicate dorsal, while positive bending angle values indicate ventral. n=3.

(G) Top, confocal images of DA/DB motor neurons from control and *dve-1* mutants expressing *punc-129::*Chrimson*::SL2::BFP*. Scale bar, 5 µm. Bottom, average fluorescence intensity of DA/DB neuron cell bodies labeled by *punc-129::*Chrimson*::SL2::BFP*. Each dot represents a single DA/DB cell body, at least 15 animals per genotype were imaged.

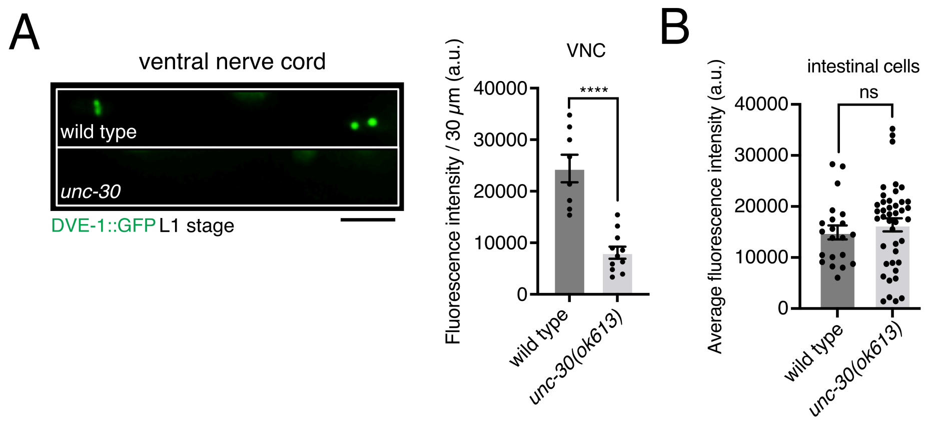


**Supplemental Figure 5.1. The Pitx family transcription factor UNC-30 regulates expression of DVE-1 in GABAergic neurons**

A. Top, confocal fluorescence images of ventral nerve cord expression of DVE-1::GFP in DD GABAergic motor neurons of L1 stage wild type and *unc-30(ok613)* mutants. Scale bar, 5 µm. Bottom, scatterplot of average fluorescence intensity measures of ventral nerve cord (VNC) 30 µm ROI. Each dot represents a single animal. wild type n=8, *unc-30(ok613)* n=11. Bars indicate mean ± SEM. ****$p$<.0001, student's t-test.

B. Scatterplot of average nuclear DVE-1::GFP fluorescence intensity in intestinal cells of L1 stage wild type and *unc-30(ok613)* mutants. Each point represents a single intestinal cell. Imaged 3 intestinal cells per animal. Wild type n=8, *unc-30* mutants n=11. Bars indicate mean ± SEM. ns - not significant, student's t-test.

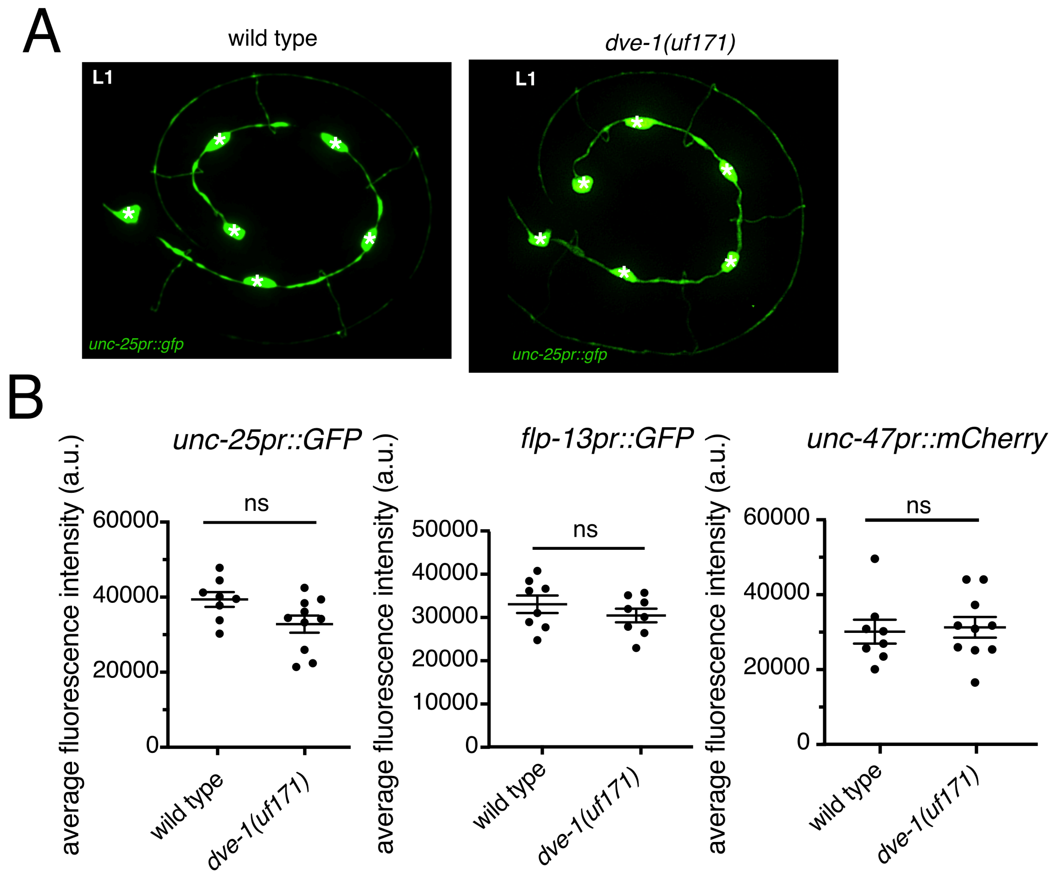


**Supplemental Figure 6.1. Mutation of *dve-1* does not alter GABAergic neuronal identity**

(A) Representative image of wild type (right) and *dve-1(uf171)* mutant (left) animals labeling DD neurons with *punc-25::GFP* at the L1 developmental stage. *-Cell body.

(B) Average fluorescence intensity of DD1, DD2, and DD3 neuron cell bodies in three distinct GABA reporter backgrounds, *unc-47pr::mCherry,* and *unc-25pr::GFP, flp-13pr::GFP* in wild type and *dve-1(uf171)* mutants at the L1 developmental stage. Each dot represents the average of three cell bodies in a single animal, ≥8 animals per genotype. Bars indicate mean ± SEM. ns-not significant, students t-test.

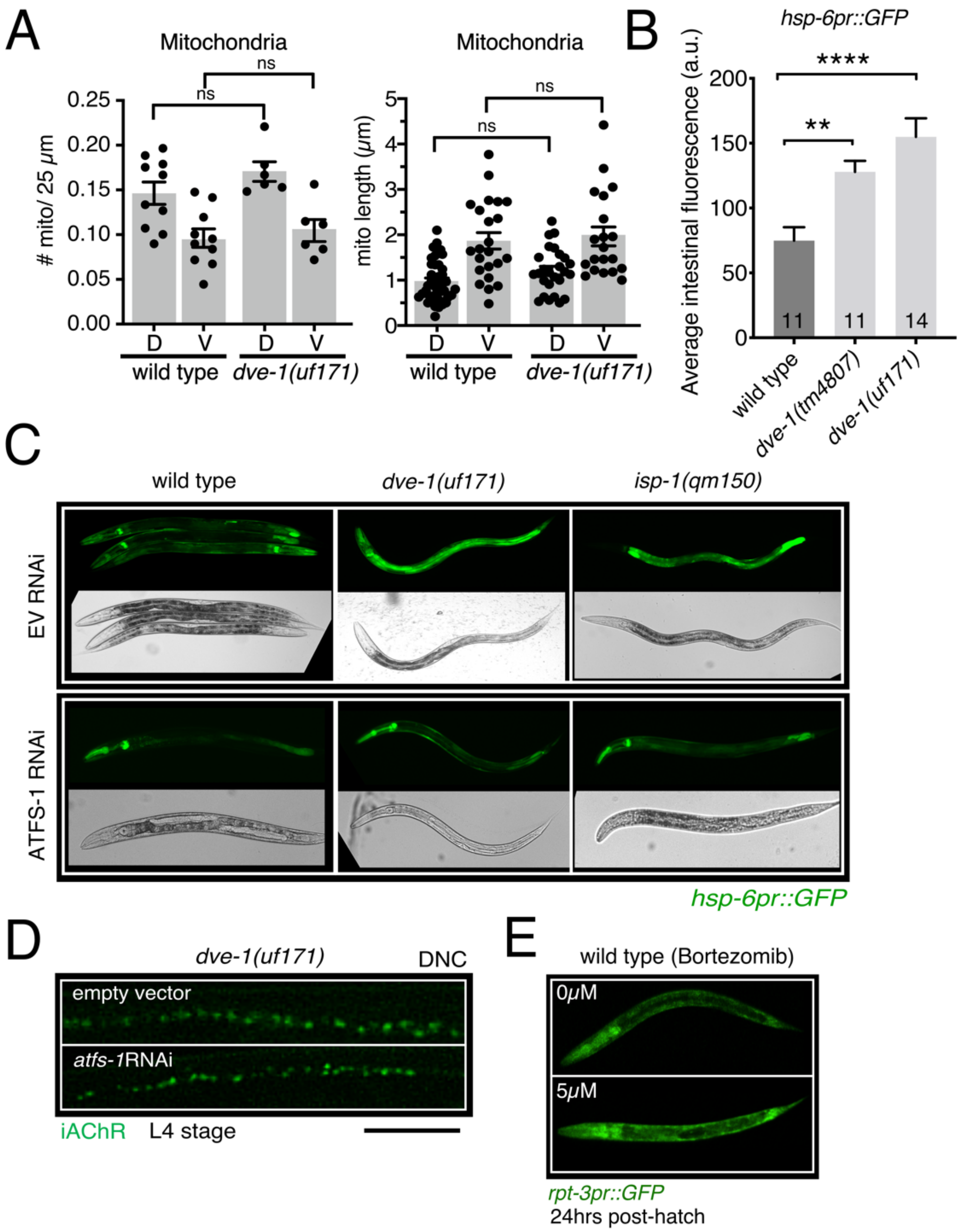


**Supplemental Figure 6.2. Removal of iAChRs in GABA MNs is not disrupted by either activation or inhibition of the mtUPR**

(A) Left, scatterplot of mitochondrial density (number of mitochondria/25 μm) in dorsal (D) and ventral (V) processes of DD neurons of L4 stage wild type and *dve-1(uf171)* mutants. Each point represents a single animal, wild type n=10, dve-1(uf171) n=6. Right, average length of mitochondria in dorsal and ventral processes of DD neurons of L4 stage wild type and *dve-*

*1(uf171)* mutants. Each dot represents a single mitochondria, wild type n=10, *dve-1(uf171)* n=6. Bars indicate mean ± SEM, ANOVA with Dunnett's multiple comparisons test. ns- not significant.

(B) Average fluorescence intensity measures of intestinal ROI in wild type, dve-1(tm4807), and dve-1(uf171) animals expressing *hsp-6pr::GFP* transgene. Bars represent mean ± SEM. ANOVA with Dunnett's multiple comparisons test. ** $p < 0.01$, **** $p < 0.0001$.

(C) Fluorescent images of transgenic worms expressing the mtUPR reporter *hsp-6pr::*GFP and treated with either empty vector (top) or RNAi targeting *atfs-1* (bottom). *dve-1(uf171)* mutants show increased expression of *hsp-6pr::*GFP under basal conditions compared with control animals and this is reversed by RNAi targeting *atfs-1*. *isp-1* mutants also have elevated mtUPR and are included as a control.

(D) Fluorescent confocal images of iAChR clusters in dorsal nerve cord (DNC) of DD neurons of L4 stage *dve-1(uf171)* mutants treated with either empty vector or RNAi targeting *atfs-1*. *atfs-1* RNAi reverses the elevated mtUPR of *dve-1* mutants but does not normalize synapse elimination. Scale bar, 5 μm.

(E) Fluorescent images of transgenic worms expressing the reporter *rpt-3pr::GFP* with and without treatment of the proteasome inhibitor Bortezomib (5 μM).
